# Decline of DNA damage response along with myogenic differentiation

**DOI:** 10.1101/2023.06.27.546688

**Authors:** Haser Hasan Sutcu, Phoebe Rassinoux, Lise-Marie Donnio, Damien Neuillet, François Vianna, Olivier Gabillot, Pierre-Olivier Mari, Céline Baldeyron, Giuseppina Giglia-Mari

**Author notes:** These authors contributed equally to the work (Céline Baldeyron, Giuseppina Giglia-Mari). Corresponding authors (Giuseppina Giglia-Mari, Céline Baldeyron**) Email:**. **Author Contributions:** HHS designed, planned, performed experiments and analyzed, processed the data. HHS also contributed to writing of the manuscript. FV supervised and performed α-particle irradiation experiments. PR, LMD, DN and OG performed laser irradiation experiments. PR also contributed to manuscript preparation. POM performed experiments and analyzed the data. CB and GGM supervised the study, analyzed the data, and wrote the manuscript. All authors read and approved the final manuscript. **Competing Interest Statement:** No competing interests. **Classification:** BIOLOGICAL SCIENCES – Cell Biology.

## Abstract

DNA integrity is incessantly confronted to endogenous and exogenous agents inducing DNA lesions, which are harmful for cellular homeostasis. Luckily all organisms are equipped with a network of DNA damage response (DDR) mechanisms that will repair DNA lesions and restore the proper cellular activities. Despite DNA repair mechanisms have been revealed in vitro and in replicating cells, still little is known on how DNA lesions are repaired and consequently how cellular homeostasis is maintained in post-mitotic cells. Muscle fibers are highly specialised post-mitotic cells organized in syncytia and, they are vulnerable to age-related degeneration and atrophy following radiotherapy treatment. We have here studied in detail the DNA repair capacity of muscle fibers nuclei and compared it with the one measured in proliferative myoblasts. We focused on the DNA repair mechanisms that correct ionizing radiation (IR)-induced lesions, namely the base excision repair (BER), the non-homologous end joining (NHEJ) and the homologous recombination (HR). We found that in the most differentiated myogenic cells, myotubes, all of these DNA repair mechanisms present weakened kinetics of recruitment of DNA repair proteins to IR-damaged DNA. For BER and HR, this decline can be linked to reduced steady state levels of key proteins involved in these processes, probably since nuclei within muscle fibers no longer replicate their DNA.

**SIGNIFICANCE:** Although skeletal muscle tissue has been considered as resistant to injury caused by ionizing radiations (IR), in the long term, muscle surrounding irradiated regions of the body following radiotherapy has physiological consequences such as atrophy and muscle wasting. To prevent and/or counteract the harmful effects of IR in muscle fibers, it is essential, to evaluate the DNA repair capacity of post-mitotic myonuclei. By using fluorescently tagged DNA repair proteins expressed in myoblasts and myotubes, we were able to assess the kinetics of the DNA damage response in these cells and determine that DNA damage response is weakened during the process of myofibrillogenesis.

## INTRODUCTION

Proper functioning of all living organisms depends on the faithful maintenance and transmission of genomic information stored in the molecule of DNA. However, DNA integrity is continuously challenged by a variety of endogenous and exogenous agents causing DNA lesions, which have a critical impact on cellular activities and homeostasis. The biological consequences of DNA lesions are varied and mostly depend on the replicative versus post-mitotic state of the cells. While in replicative cells, the acute effects of DNA damage arise from the disturbance of DNA replication leading to irreversible mutations, in non-replicative post-mitotic cells, DNA lesions physically block transcription causing general cellular dysfunction and premature cell death ^1^. In order to prevent the deleterious consequences of persisting DNA lesions, all organisms are equipped with an intricate network of DNA damage response (DDR) mechanisms ^1, 2^ covering most of the genomic insults. While DNA Repair mechanisms have been thoroughly described *in vitro* and in replicating cells, but little is known on these processes and their role in the maintenance of the cellular homeostasis in post-mitotic cells.

Post-mitotic cells represent the majority of cells in our adult body and amongst them, <u>S</u>keletal <u>M</u>uscle <u>F</u>ibers (SMFs) represent almost 40% of the body mass ^3^. SMFs are highly specialised post-mitotic cells organized in syncytia resulting from the fusion of hundreds of myoblasts ^3^. Before fusion, myoblasts are highly proliferative, then they exit the cell cycle and become myocytes possessing the potential to fuse with each other. Homeostasis of the adult muscle is insured by <u>M</u>uscle <u>S</u>tem <u>C</u>ells (MuSCs), also named satellite cells ^3^, which lay quiescent in their niche along the myofiber, under the basal lamina ^3^. MuSCs can be isolated from muscles and their proliferation and differentiation can be achieved and scrutinized *in vitro* ^3^. Each myonucleus within SMFs has to deal with 10^4^-10^5^ lesions per day and despite the ability of SMFs to partially regenerate, muscle fibers age with the organism and have to deal with this damage load, making them vulnerable to degeneration from age-related disturbances in cellular homeostasis^4^. In fact, muscle cachexia and atrophy are observed in many physiological, traumatic and pathological situations ^4^. A classical DNA damage-induced muscular atrophy is observed after radiotherapy treatment. In fact, although the skeletal muscle tissue has been considered as radio-resistant ^5–7^, several studies show that, in the long term, irradiation has physiological consequences on the muscle depending on the dose, frequency, or type of radiation ^8, 9^. These complications include muscle wasting, cachexia, contractures, malfunctioning and weakness, and can even be more severe for the juvenile patients who are still under development ^5, 10, 11^. Ionizing radiations (IR) induces a plethora of different types of damage, ranging from base damages, abasic sites, oxidation of bases, single strand breaks (SSBs) repaired via the <u>B</u>ase <u>E</u>xcision <u>R</u>epair (BER) and SSBR pathways ^12^, which converge in the same path in the final steps, and double strand breaks (DSBs) repaired by <u>N</u>on-<u>H</u>omologous <u>E</u>nd-<u>J</u>oining (NHEJ) in post-mitotic cells ^1^. BER consists of 2 sub-pathways: short-patch and long-patch BER. BER is initiated by specific DNA Glycosylases dependent recognition and removal of a damaged base, then under coordination of PARP1, DNA is cleaved by AP endonuclease 1 (APE1) ^13, 14^. In short patch BER, a correct nucleotide is incorporated, and ligation of nicked DNA ends the repair reaction ligated by the complex XRCC1/Ligase 1 or Ligase 3. During long-patch BER, AP endonuclease 2 (APE2) provides longer resection, and 2-12 nucleotides are incorporated to the DNA damage site, which is then further processed by the Flap Structure-Specific Endonuclease 1 (FEN1) ^15^ and finally ligated ^14, 16^. In post-mitotic cells, repair of DSBs is insured by the NHEJ, initiated by recruitment of KU70-KU80 heterodimer ^17^, followed by the DNA-dependent protein kinase catalytic subunit (DNA-PKcs) allowing the broken DNA ends to be processed and, subsequently, ligated by Ligase 4 ^18^ along with its mediators XRCC4 and XLF/Cernunnos ^19, 20^.

In SMFs, previous work has shown that levels of oxidative damage are increased compared to myoblasts and that BER is attenuated ^21^. It has also been reported that DSBs repair efficiency is increased in MuSCs compared to committed progenitors ^22^. These studies show that there is indeed a difference in the DNA repair activity between MuSCs and SMFs but remain anecdotical and a more in-depth investigation is needed to disclose whether differences in DNA repair activity effectively exist during myofibrillogenesis.

Here we performed a systemic study to assess and increase our understanding in DNA damage repair mechanisms specific to different stages of myogenesis from mono-nuclear precursor cells until fused multi-nuclear myotubes. By using myoblasts isolated from a fluorescently tagged Fen1 knock in mouse model ^15^ and, immortalized and primary myoblasts expressing fluorescently tagged DNA damage signaling and repair proteins we were able to assess the kinetics of DDR during the process of myofibrillogenesis.

## RESULTS

### Transcriptional activity by the RNAP1 and RNAP2 during myofibrillogenesis

The most abundant DNA lesions induced by ionizing radiation (IR) treatment are oxidatively damaged bases and single strand breaks (SSBs) ^23^, which are repaired by Base Excision Repair (BER) pathway. In order to study Base Excision Repair (BER) activity of myoblasts versus myotubes, we isolated myoblasts from muscles of 5-days old mice from the mouse models expressing endogenously a fluorescent tagged version of Fen1 ^15^ and differentiate them to a full myotube syncytium (Fig S1). As a first step in our study, to identify which key steps during myofibrillogenesis had to be investigated, we decided to examine how transcriptional activity is modified during myofibrillogenesis. In fact, it has been shown that in post-mitotic cells, DNA repair pathways act mainly on transcribed regions of the genome ^24, 25^ and we wanted to verify that, during myofibrillogenesis, the general transcriptional activity was not dissimilar, which could have explained differences in DNA repair activities. We chose to measure both RNA Polymerase 2 (RNAP2) and RNA Polymerase 1 (RNAP1) activity as previously described ^26, 27^ and we selected 4 different steps of the differentiation, namely: (i) myoblasts, (ii) myocytes in fusion, (iii) myotubes at 4 days of differentiation and (iv) myotubes at 7 days of differentiation. RNAP2 activity was measured using 5-Ethinyl-Uridine (EU) incorporation into newly synthetised mRNA (Fig S2A). We detected a RNAP2 transcriptional activity increased by a 3-fold change in fusing myocytes compared to the one in myoblasts, while myoblasts and myotubes at 4 or 7 days of differentiation have a more similar, but still statistically different, RNAP2 transcriptional activity (Fig S2B). RNAP1 activity was measured using an RNA-FISH assay, specifically labelling the 47S pre-ribosomal RNA species (Fig S2C and S2D). RNAP1 activity does not increase in myoblasts in fusion, as RNAP2, and does not follow the change of the RNAP2 activity during myofibrillogenesis it mainly decreases slowly during differentiation (Fig S2E). These results led us to further study the DNA repair activity of proliferative myoblasts and compare it to the one measured in 7 days differentiated myotubes.

### Myotubes have weakened Base Excision Repair competence than myoblasts

In order to measure the Base Excision Repair (BER) activity in myoblasts versus myonuclei in myotubes, we isolated myoblasts from 5-day old pups. At this age, muscles are continuously growing and satellite cells, which in the adult muscles are quiescent, are highly proliferative and have myoblasts properties ^28^. To induce local oxidative base damage, we used different approaches: (i) multiphoton laser-beam damage induction and (ii) a targeted α-particle irradiation by using a focused heavy ion microbeam. Multiphoton damage is obtained with near-infrared tuneable laser (Coherent). This type of localised laser-irradiation induces a plethora of different DNA lesions, amongst which oxidative damage, without the addition of DNA intercalators that could induce chromatin disturbances and affect different cellular activities ^29^. To be able to measure just the BER activity, we isolated myoblasts from a mouse model that endogenously express the specific BER protein FEN1, here after FEN1-YFP ^15^. To verify that the accumulation of Fen1-YFP on the damaged substrate is indeed due to the DNA repair process and in response to DDR, we performed the assay (schematic representation for quantification of fluorescent-tagged protein recruitment to local damage, Fig S3) in presence of different DDR inhibiting drugs in fibroblasts isolated from FEN1-YFP mouse models (Fig S4A). Without any DDR inhibiting drugs FEN1-YFP is rapidly recruited to the damaged DNA and is progressively released from the damage as the BER process advances (Fig S3A), however FEN1-YFP recruitment is partially impaired in presence of KU55993 (ATM inhibitor ^30^ and VE821 (ATM/ATR inhibitor ^31^(Fig S4A). The recruitment of FEN1-YFP is even more diminished when cells are treated with both inhibitors at the same time (Fig S4A). Our results are thus in agreement with previously published data showing that ATM- and ATR-dependent checkpoint pathways are required to coordinate DNA repair process in the presence of oxidatively damaged DNA ^32, 33^.

We performed the same assay in myoblasts (MB) and myonuclei within myotubes (MT) and interestingly, we could observe that the BER repair kinetics are different in MB versus MT. In fact, while MB repair kinetics are very similar to the ones measured in fibroblasts (Fig 1A), MT have a reduced recruitment and a slower repair kinetics, indicating that more than half of the BER substrate is still present 30 minutes after damage induction (Fig 1A). This unexpected result prompted us to explore whether the different repair kinetics is related to the fact that myonuclei are in a syncytium or if it is an intrinsic characteristic of differentiated myotubes. In order to verify this hypothesis, we have performed the same measurements of DNA repair kinetics by laser-damage induction within fibroblasts that have been forced to create a syncytium. The results, presented in (Fig 1B), show that fused fibroblasts present a reduced recruitment of FEN1-YFP but a fast release from the substrate. Because MT are post-mitotic cells and do not need FEN1 for its replication function, we wondered whether the FEN1 steady state concentration would impact the level of FEN1 recruitment on the local DNA damage (LD) induced by laser-irradiation. In order to establish a correlation between these two parameters, we measured the steady state concentration of FEN1-YFP and compared the correspondent maximum level of recruitment (Fig S5A). The recruitment level of FEN1-YFP in both fused fibroblasts and MT correlates with the steady state level of FEN1-YFP protein in these cells (Fig S5B), suggesting that, in these cells, FEN1 could be rate-limiting for the BER reaction. However, despite a reduced recruitment, the release from the damaged substrate, which is a direct measure of the DNA repair activity of the cells, in fused fibroblasts are comparable to the ones measured in fibroblasts and MB (Fig 1A and 1B and Fig S5). In summary, the ½ life of substrate (oxidative lesions) in MB, fibroblasts or fused fibroblasts is in the range of 400 to 700 sec, while the ½ life of the substrate in MT is not yet reached between 1000 and 1200 sec (Fig S6). The reduction of DNA repair activity is just observed in differentiated myonuclei within myotubes (Fig 1A).

**Figure 1.**
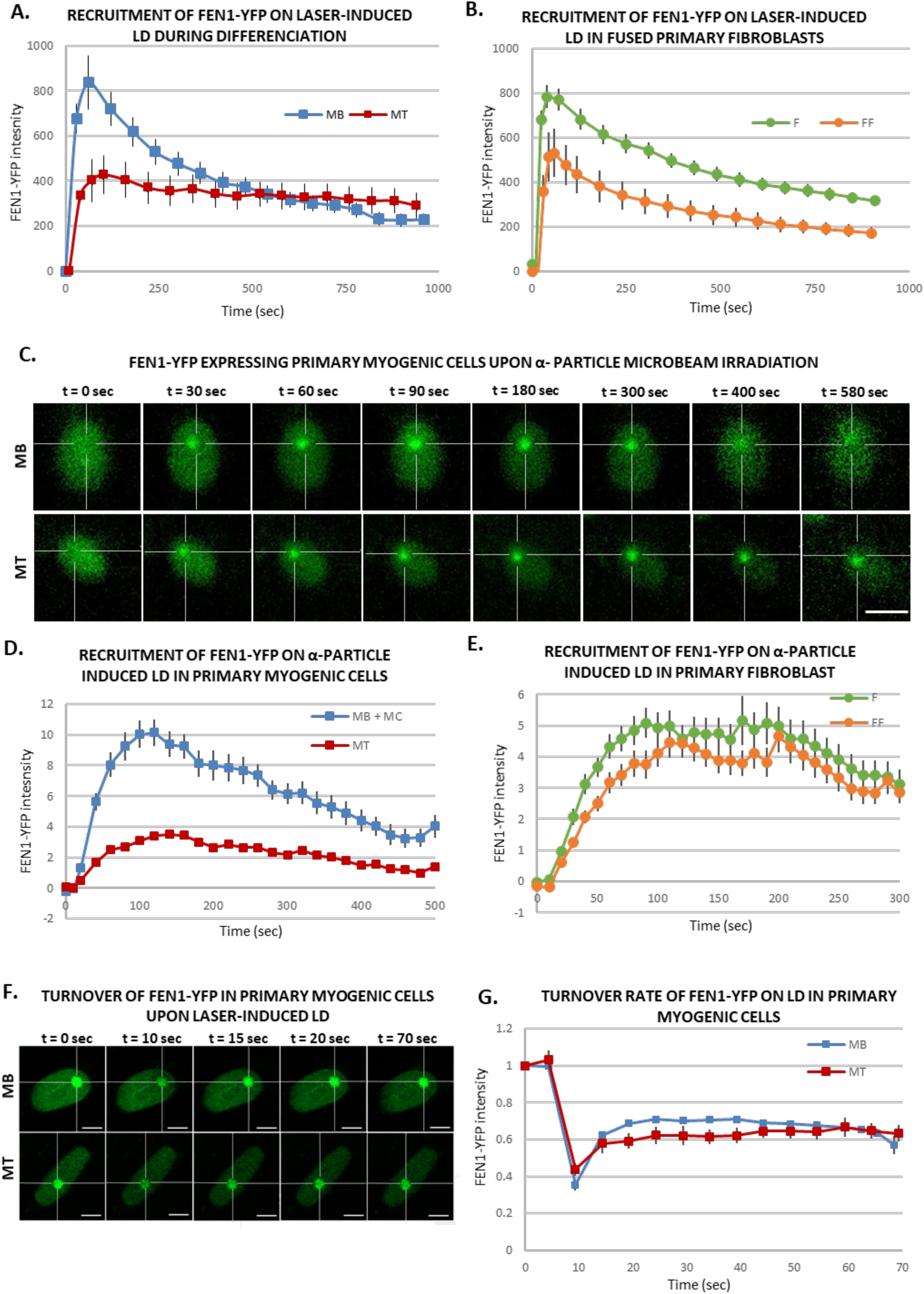
Base Excision Repair (BER) activity during myofibrillogenesis. **A, B.** Recruitment curve of FEN1-YFP on the locally damaged DNA (LD) by laser-micro irradiation (A) in myoblasts (MB, blue curve) isolated from the Fen1-YFP mouse model) and 7 days differentiated myonuclei (MT, red curve) and (B) in primary dermal fibroblast (F) isolated from the Fen1-YFP mouse model (green curve) and the same fibroblasts fused with PEG (FF, orange curve). Error bars represent the SEM obtained from at least 15 nuclei in (A) and, 19 nuclei for F and 12 nuclei for FF cells in (B) from N≥3 independent experiments. **C.** Sequential images of FEN1-YFP recruitment onto LD by α-particle microbeam irradiation in MB (upper panel) isolated from the Fen1-YFP mouse model and in MT (lower panel). The damaged areas are underlined by a dotted cross within the nucleus. The scale bar represents 10 µm. **D, E.** Recruitment curve of FEN1-YFP on the LD by α-particle microbeam irradiation (D) in MB (blue curve) and MT (red curve) and (E) in primary dermal fibroblast (F) isolated from the Fen1-YFP mouse model (green curve) and the same fibroblasts fused with PEG (FF, orange curve). The irradiation was applied at t = 10 s. Error bars represent the SEM obtained from N≥3 independent experiments with 42-46 nuclei/cell type in (D) and 37-47 nuclei/cell type in (E). **F.** Sequential imaging of FEN1-YFP turnover on the LD by laser-irradiation in MB (upper panel) and MT (lower panel). The damaged area is underlined by a dotted cross in the nucleus. Scale bar, 5 µm. **G.** Turnover curve of FEN1-YFP on the LD by laser-irradiation in MB (blue curve) and MT (red curve). Error bars represent the SEM obtained from from N≥3 independent experiment with 10 nuclei.

The laser microbeams play a major role in the study of the temporal and spatial organization of the cellular DNA damage response, by allowing the induction of DNA damage in a defined region in the cell nucleus *in situ* with a micrometric precision and permitting the monitoring of recruitment kinetics of DNA damage response (DDR) proteins to localized DNA damage sites ^34^. However, the heavy ion microbeam technology offers in addition the possibility to deliver a predetermined number of particles of a certain radiation quality (type and energy) ^35^ . This technology was used to investigate whether this reduction of repair of oxidatively damaged DNA is a general behaviour upon treatment with other genotoxic agents. Irradiation with α-particles is known to induces, in addition to DNA strand breaks, oxidative base lesions ^36^. We thus performed locally irradiation within cell nuclei with a predetermined number of 6 MeV α-particles with a micrometric spatial resolution ^37, 38^. We measured the BER activity in mononuclear cells including MB and myocytes (MB+MC), which are mononuclear myogenic cells, and in MT upon local irradiation with 1,000 α-particles (Fig 1C and 1D). We also found that the BER repair kinetics are different in MB+MC versus MT (Fig 1D) and, MT have a reduced recruitment and a slower repair kinetics (Fig 1D). We have carried out the same measurements within fused fibroblasts. As with laser-irradiation, fused fibroblasts present after local α-particles irradiation a reduced recruitment of FEN1-YFP and a release from the substrate as fast as in fibroblasts (Fig 1E). Together the data obtained with local α-particles irradiation confirmed that the decrease of BER activity is a characteristic of differentiated myonuclei (Fig 1C and 1D). We wondered whether this difference could be due to a deficient turnover of FEN1 due to a different upstream and downstream binding partners’ level. We measured this turnover rate by using FRAP on LD (Fluorescence Recovery After Photobleaching on Local Damage) (Fig 1F) and we could measure no difference in FEN1 turnover between MB and MT (Fig 1G). To investigate whether the decrease in BER activity could be due to a decrease in the steady state level of BER related proteins, we performed immunofluorescences (IF) and observed that FEN1, LIG1, XRCC1, PARP1, APE1 are all downregulated in MT compared to MB (Fig S7A). All together these results point to a deficiency in BER activity in MT compared to MB probably due to an overall decreased steady state level of BER proteins.

### Double Strand Break repair in myotubes is weaker than in proliferative myoblasts

While few DNA double-strand breaks (DSB) are produced upon irradiation, DSB is the most critical lesion, which when mis-repaired or unrepaired, can lead to genomic instability and cell death ^39^. Previously, it was described that skeletal muscle stem cells repair ionizing radiation (IR)-induced DSBs more efficient than their committed progeny ^22^. To clearly assess the differences in DSB repair efficiency between myoblasts (MB) and myonuclei within myotubes (MT), C2C7 ^40^(immortalized murine myoblast cell line) MB were differentiated into myocytes (MC) and myotubes (MT) and irradiated with 5 Gy of X-ray using medical linear accelerator (LINAC, Elekta Synergy®). As we performed this kind of assay in C2C7 cells stably expressing GFP-tagged 53BP1 (Fig 2), we first confirmed that the behavior of 53BP1-GFP is similar to this of endogenous 53BP1. Upon irradiation (Irr), cells were kept in culture and examined at different time points post-Irr (i.e. 2 hours to 2 days) in order to quantify γ-H2AX and 53BP1 IR-induced foci (IRIF) ^41, 42^ as a measure of DSBs presence/signaling and repair. In C2C7 cells transiently transfected with 53BP1-GFP plasmids, irradiation with 5 Gy of X-ray induced the formation of 53BP1-GFP and endogenous 53BP1 foci (Fig S8A) that disappear with the same kinetics (Fig S8B). We found that the exogenous expression of 53BP1 in C2C7 MB has no impact on the appearance and disappearance of γ-H2AX foci upon 5 Gy of X-ray Irr (Fig S8C). By performing the same type of experiment, we thus quantified the γ-H2AX and 53BP1 IRIF in myogenic cells stably expressing 53BP1-GFP (Fig 2). At early time points after Irr (2 hours post-Irr), the presence of DSBs was confirmed by the increased number of γ-H2AX and 53BP1-GFP foci in MB, MC and MT (Fig 2A and 2B). Interestingly, MT had lower number γ-H2AX foci (Fig 2A and 2B) and 53BP1-GFP foci (Fig 2A and 2C) in comparison to MB. Interestingly, while in proliferative MB, at 1 day post-Irr, both γ-H2AX and 53BP1-GFP foci numbers were significantly decreased to reach similar levels as the non-irradiated (Non-Irr) condition, MT showed some decrease in number of foci, although they remained higher in comparison to MB, indicating still presence of DSBs at 24 hours post-Irr (Fig 2B and 2C).

**Figure 2.**
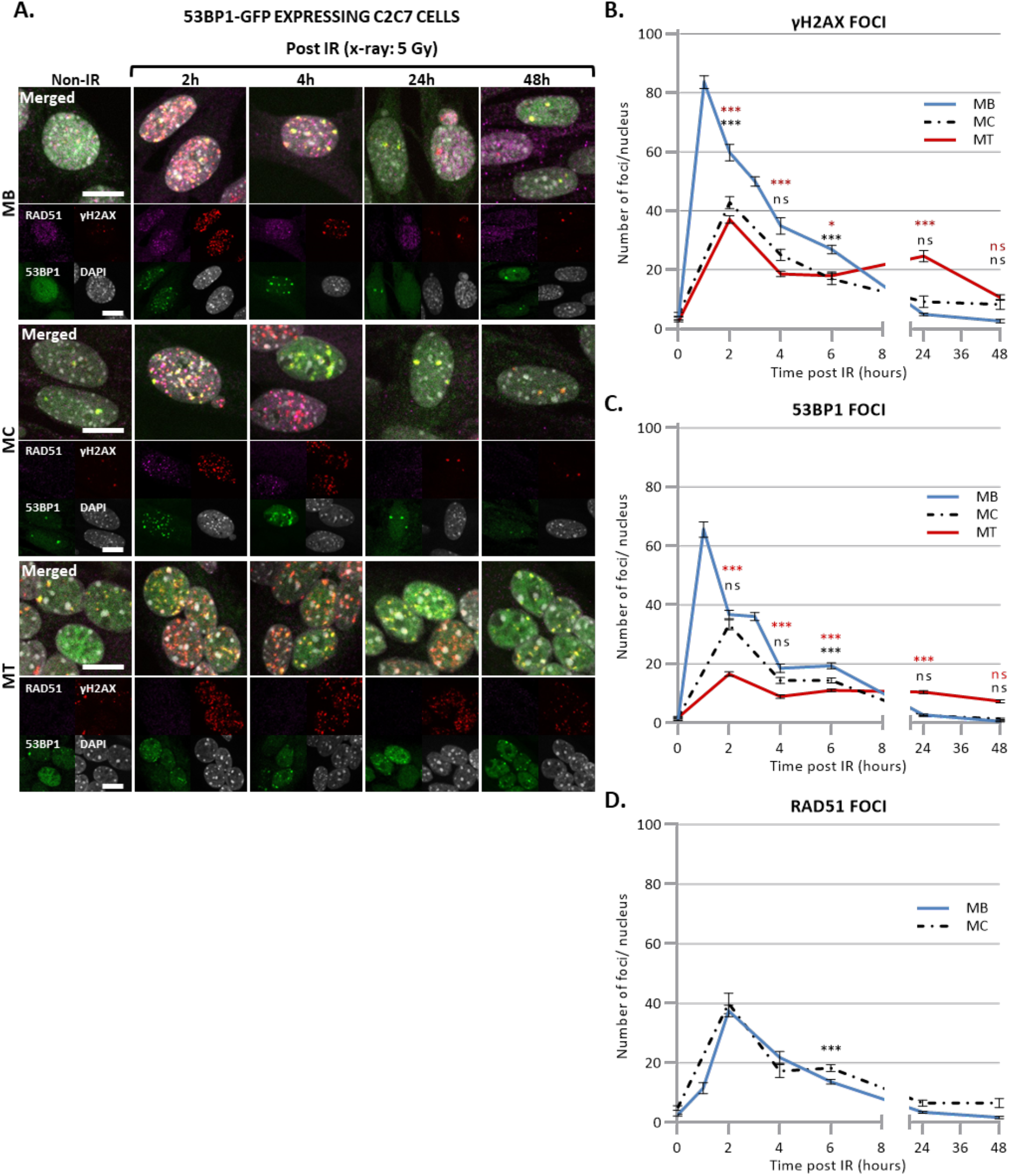
Disappearance of DSB signaling upon X-ray irradiation. **A.** Representative images of stably 53BP1-GFP (green) expressing C2C7 myogenic cells at different state of differentiation, myoblasts (MB, upper panel), myocytes (MC, middle panel) and myotubes (MT, lower panel) at the indicated time post-5 Gy of X-ray irradiation, immunolabelled with antibodies against the homologous recombination (HR) factor RAD51 (magenta), and a DSB marker, the phosphorylated histone H2AX protein (γ-H2AX, red). DNA was stained with DAPI (grey). Scale bars, 10 µm. **B, C, D.** Quantification of γH2AX foci (B), 53BP1 foci (C) and RAD51 foci (D) per nucleus in stably 53BP1-GFP expressing C2C7 MB (blue curve), MC (dashed black curve) and MT (red curve) upon 5 Gy of X-ray irradiation. N ≥ 3 independent experiments with 35-135 nuclei/cell type. Mean ± SEM, significance by 1-way ANOVA with post-hoc Tukey’s multiple comparison test against MB at each time point, significant p-value figures are same colour as the condition compared ^ns^ p>0,05, * p<0.05, ** p<0.01, *** p<0.001.

Taken together, these results strongly suggest that DSBs signaling and/or repair is impaired or reduced in MT, compared to MB and MC.

### Double Strand Breaks in myotubes are not repaired by HR

DSBs are repaired by either HR or NHEJ ^43, 44^. Unlike NHEJ that operates at all stages of the cell cycle in replicative cells, HR is restricted to S and G2 phases of cell cycle when the homology donor is nearby. Thus, in post-mitotic cells the DSB repair pathway of choice is the NHEJ ^45, 46^. The key protein of DSB repair mediated by HR is RAD51^47^, which plays a fundamental role in mediating invasion of homologous template DNA ^48^. Predictably, in irradiated post-mitotic MT stably expressing 53BP1-GFP, we could not observe any RAD51 foci, validating the absence of HR (Fig 2A and 2D) in post-mitotic cells ^49^. Interestingly, no RAD51 was detectable by IF in MT, suggesting that MT have either no or indetectable expression of RAD51 (Fig. S7B).

These results suggest that MT have a declined DSBs repair by the HR machinery and that DSBs in these post-mitotic cells are likely to be exclusively repaired by NHEJ.

### Double Strand Breaks in myotubes are repaired by a weakened NHEJ

To investigate the dynamic of NHEJ during myofibrillogenesis, we produced a C2C7 cell line stably expressing KU80-GFP and assessed the recruitment capacity of this NHEJ factor to the induced local DNA damage site at different steps of myofibrillogenesis. KU70/80 is an heterodimer essential for the detection and repair of DSBs during NHEJ ^17^, in this pathway KU70/80 is recruited to the damaged DNA ends, protecting them from nuclease activity and being a platform for the subsequent steps of NHEJ ^50^. We induced in nuclei of KU80-GFP stably expressing cells local DSBs by using a near-infrared multiphoton laser (which was also previously used to study the dynamic assembly of NHEJ factors ^17^) (Fig 3A and 3B) and α-particles microbeam (Fig 3C and 3D) in both MB and MT and follow the recruitment of KU80-GFP over a time frame of several minutes (5 minutes for the α-particles damage and 10 min for the laser damage). Using these damage induction systems, we observed a clear difference in the recruitment of KU80-GFP on the damaged DNA in MB versus MT (Fig 3B and 3D). We obtained similar results when we assessed the KU80 kinetics in primary isolated myoblasts transiently transfected with KU80-GFP, a weaker KU80-GFP recruitment in MT (Fig S9A and S9B). In fact, while replicating MB showed a repair kinetics very similar to the one previously observed in KU80 complemented CHO cells ^17^, post-mitotic MT presented a reduced KU80-GFP recruitment (approximately half of the KU80-GFP recruitment measured in MB). As for the reduced recruitment of FEN1-YFP in MT (Fig 1), the low recruitment of KU80 on damaged DNA in MT could be explained by the difference in steady state levels of KU80 in MT versus MB, however unlike FEN1, the amount of KU70/80 heterodimer was a bit higher in MT when compared to MB (Fig S7C). Another plausible explanation would be that on damaged DNA KU80 turnover is faster in MT (compared to Ku80 turnover in MB), implying a reduced occupancy of the damaged substrate. We could confirm this hypothesis by performing Fluorescence Recovery After Photobleaching on Local Damage (FRAP on LD) in MB and MT (Fig 3E). Using this FRAP variation, we could estimate the turnover rate of KU80-GFP on damaged DNA after 10 min of damage induction and show that KU80 is rapidly exchanging with the damaged substrate in post-mitotic MT while it has a slower turnover rate in replicative MB, showing that in these latter cells, KU80-GFP is more strongly bound to the substrate (Fig 3F). These results might indicate that in MT stabilizing factors maintaining KU80 on the DNA ends might be under expressed or not functional. To confirm that NHEJ process was also impacted at the late steps, we measured that dynamic of recruitment and repair of LIG4, the ATP-dependent DNA ligase responsible for ligation of the broken DNA ends during NHEJ ^51^. We performed laser-damage (Fig 4A and 4B) and local α-particles irradiation in LIG4-GFP stably expressing C2C7 cells (Fig 4C and 4D) and LIG4-GFP transiently transfected primary myoblasts, and observed that in all cases the recruitment of DNA Ligase 4 was reduced in MT compared to MB (Fig 4A, 4B, 4C and 4D and Fig S9C and S9D). We confirmed that this highly decreased accumulation of LIG4-GFP at the local site of Irr-damaged DNA was not due to a limit amount of this protein (Fig S7D). In addition, we performed FRAP on LD to measure the turnover of LIG4 and demonstrate that there is no change in the turnover rate of this protein on the LD (Fig 4E and 4F)

**Figure 3.**
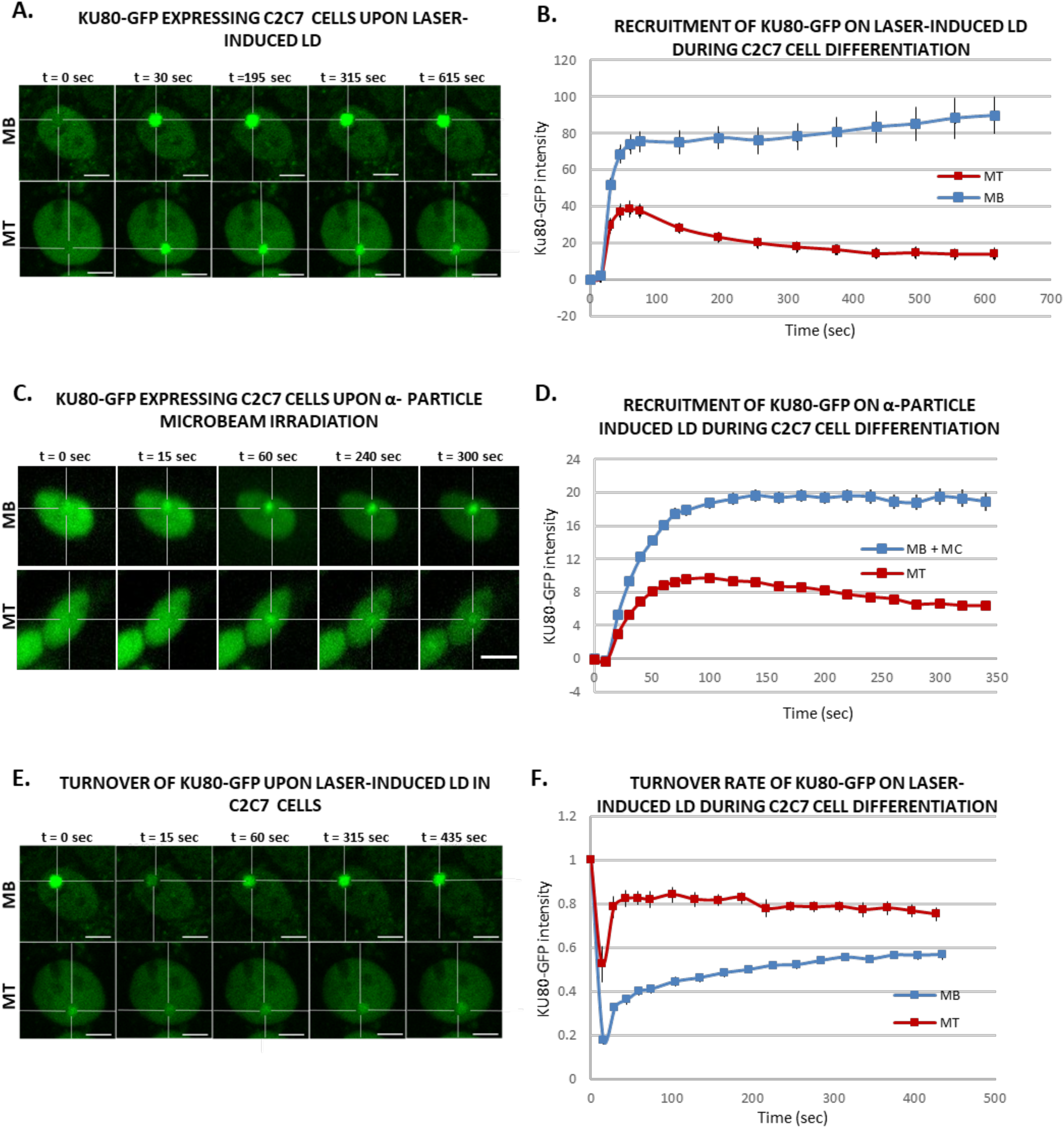
Activity of an early Non-Homologous End-Joining (NHEJ) protein during myofibrillogenesis. **A.** Sequential imaging of KU80-GFP recruitment onto the locally damaged DNA (LD) by laser-micro irradiation in stably KU80-GFP expressing C2C7 myoblasts (MB, upper panel) and myotubes (MT, lower panel). The damaged area is underlined by a dotted cross in the nucleus. Scale bar, 5 µm. **B.** Recruitment curve of KU80-GFP on the LD by laser-irradiation in stably KU80-GFP expressing C2C7 MB (blue curve) and MT (red curve). Error bars represent the SEM obtained from N≥3 independent experiment with 10 nuclei. . **C.** Sequential imaging of KU80-GFP recruitment onto the LD by α-particle microbeam irradiation in stably KU80-GFP expressing C2C7 MB (upper panel) and MT (lower panel). The damaged area is underlined by a dotted cross in the nucleus. Scale bar, 10 µm. **D.** Recruitment curve of KU80-GFP on the LD by α-particle microbeam irradiation in stably KU80-GFP expressing C2C7 myoblasts and myocytes (MB+MC, blue curve) and MT (red curve). The irradiation was applied at t = 10 s. N ≥ 3 independent experiments with 143-360 nuclei/cell type, and mean ± SEM. **E.** Sequential imaging of KU80-GFP turnover on the LD by laser-irradiation in stably KU80-GFP expressing C2C7 MB (upper panel) and MT (lower panel). The damaged area is underlined by a dotted cross in the nucleus. Scale bar, 5 µm. **F.** Turnover curve of KU80-GFP on the LD by laser-irradiation in stably KU80-GFP expressing C2C7 MB (blue curve) and MT (red curve). Error bars represent the SEM obtained from N≥3 independent experiment with 10 nuclei. .

**Figure 4.**
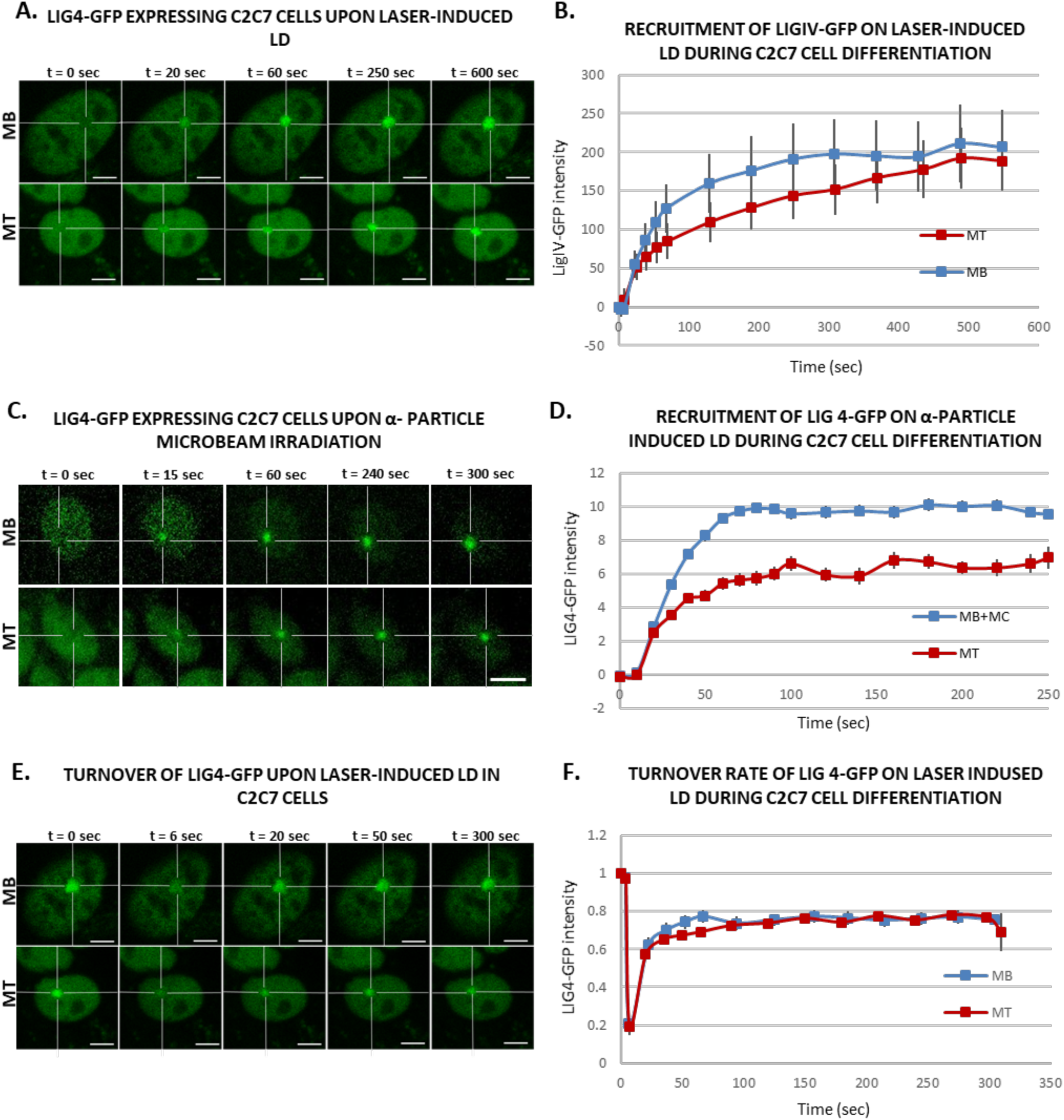
Activity of a late NHEJ protein during myofibrillogenesis. **A.** Sequential imaging of LIG4-GFP recruitment onto the locally damaged DNA (LD) by laser-irradiation in stably LIG4-GFP expressing C2C7 myoblasts (MB, upper panel) and myotubes (MT, lower panel). The damaged area is underlined by a dotted cross in the nucleus. Scale bar, 5 µm. **B.** Recruitment curve of LIG4-GFP on the LD by laser-irradiation in stably LIG4-GFP expressing C2C7 MB (blue curve) and MT (red curve). Error bars represent the SEM obtained from at least N≥3 independent experiment with 10 nuclei. . **C.** Sequential imaging of LIG4-GFP recruitment to the LD by α-particle microbeam irradiation in stably LIG4-GFP expressing C2C7 MB (upper panel) and MT (lower panel). The damaged area is underlined by a dotted cross in the nucleus. Scale bar, 10 µm. **D.** Recruitment curve of LIG4-GFP on the LD by α-particle microbeam irradiation in stably LIG4-GFP expressing C2C7 myoblasts and myocytes (MB+MC, blue curve) and MT (red curve). The irradiation was applied at t = 10 s. N ≥ 3 independent experiments with 26-138 nuclei/cell type, mean ± SEM. **E.** Sequential imaging of LIG4-GFP turnover on the LD by laser-irradiation in stably LIG4-GFP expressing C2C7 MB (upper panel) and MT (lower panel). The damaged area is underlined by a dotted cross in the nucleus. Scale bar, 5 µm. **F.** Turnover curve of LIG4-GFP on the LD by laser-irradiation in stably Lig4-GFP expressing C2C7 MB (blue curve) and MT (red curve). Error bars represent the SEM obtained from at least N≥3 independent experiment with 10 nuclei. .

Thus, our data obtained with local laser-damage and α-particles irradiation confirmed that the decrease of NHEJ activity is also a characteristic of differentiated status of myonuclei.

### Reduced 53BP1 recruitment in myotubes upon induced local DNA damage

It has been shown that 53BP1 plays a pivotal role in the choice of the DSBs mechanism, namely in proliferative cells, 53BP1 promotes error-free canonical NHEJ (c-NHEJ) over HR and error-prone alternative NHEJ (alt-NHEJ), by preventing DSB end resection ^52, 53^. However, we have shown here that in post-mitotic cells HR is impeded and this result prompted us to investigate, whether in the absence of the choice between HR and c-NHEJ, 53BP1 would be increasingly recruited of DNA damage in the few minutes upon DSB induction. In order to quantify the recruitment of 53BP1 during myogenesis we used the 53BP1-GFP stably transfected C2C7 cell line. This cell line was differentiated into MC and MT and damaged with local α-particles Irr. Interestingly, along the differentiation we could observe a progressive decrease in the recruitment of 53BP1-GFP on damaged DNA (Fig 5 A, and B) and remained low in myotubes until 1-hour post-irradiation confirmed by immuno-staining of 53BP1 and γ-H2AX in stably expressing KU80-GFP cells 7 min, 15 min and 1 hour post-Irr (Fig S10 A-B). We obtained similar results when we assessed the 53BP1 kinetics in primary isolated myoblasts transiently transfected with 53BP1-GFP, a strong decrease in 53BP1-GFP accumulation at the local Irr-damaged DNA sites in MT (Fig S11 A-B). Furthermore, immunostaining of 53BP1 and γ-H2AX in primary myogenic cells and fibroblasts fused or non-fused, isolated from FEN1-YFP mouse, 20 min post α-particles Irr, suggested that reduced recruitment of 53BP1 to the DNA damage site is specific to MT (Fig S11 C). Additionally, we observed that 53BP1 response progressively reduced with advancement of myogenic differentiation (Fig S12 A), which was also similar behavior for KU80-GFP (Fig S12 B). Despite lack of initial 53BP1 response to induced DNA damage in myotubes (Fig 5), we could observe the induced DNA damage by presence of TUNEL assay and γ-H2AX signal at the site of DNA lesions 30 min post α-particle Irr (Fig S10 C). Moreover, 6 hours and 24 hours post α-particle Irr, we have observed a late 53BP1 recruitment to the DNA damage site confirmed by presence of γ-H2AX in myotubes with no apoptosis assessed by TUNEL assay (Fig S10 C). Moreover, nuclei in myotube had much stronger γ-H2AX in comparison to myoblasts 24 hours post-Ir (Fig S10 C), confirming the slow DSB repair in myotubes observed in figure 2. To corroborate whether the different repair kinetics is an intrinsic characteristic of differentiated myotubes, we have carried out the same measurements of DNA repair kinetics with laser microbeam within C2C7 MB cells transiently transfected with 53BP1-GFP which have differentiated in MT and mouse fibroblast NIH-3T3 cells, transiently transfected with 53BP1-GFP plasmids, which have been forced to create a syncytium. After local laser-irradiation we only found a clear reduction of 53BP1-GFP recruitment in MT in comparison to MB, NIH-3T3 fibroblasts fused or non-fused (Fig S13 A-B). We confirmed this decreased accumulation of 53BP1 at damaged DNA by revealing endogenous 53BP1 by immunostaining in stably KU80-GFP expressing C2C7 myogenic cells upon local laser Irr. As expected at the local DNA damage sites revealed by a TUNEL staining ^17^, in MT we found KU80-GFP but not 53BP1 (Fig S13 C). By measuring the total nuclear 53BP1 in MB, MC and MT, we observed that MT have almost 50% and 34% higher quantity of 53BP1 than found in MC and MB respectively (Fig S7 E). Accordingly, total nuclear 53BP1 quantification suggests that the reduced recruitment of 53BP1 in MC and to a larger extent in MT is not quantity dependent.

**Figure 5.**
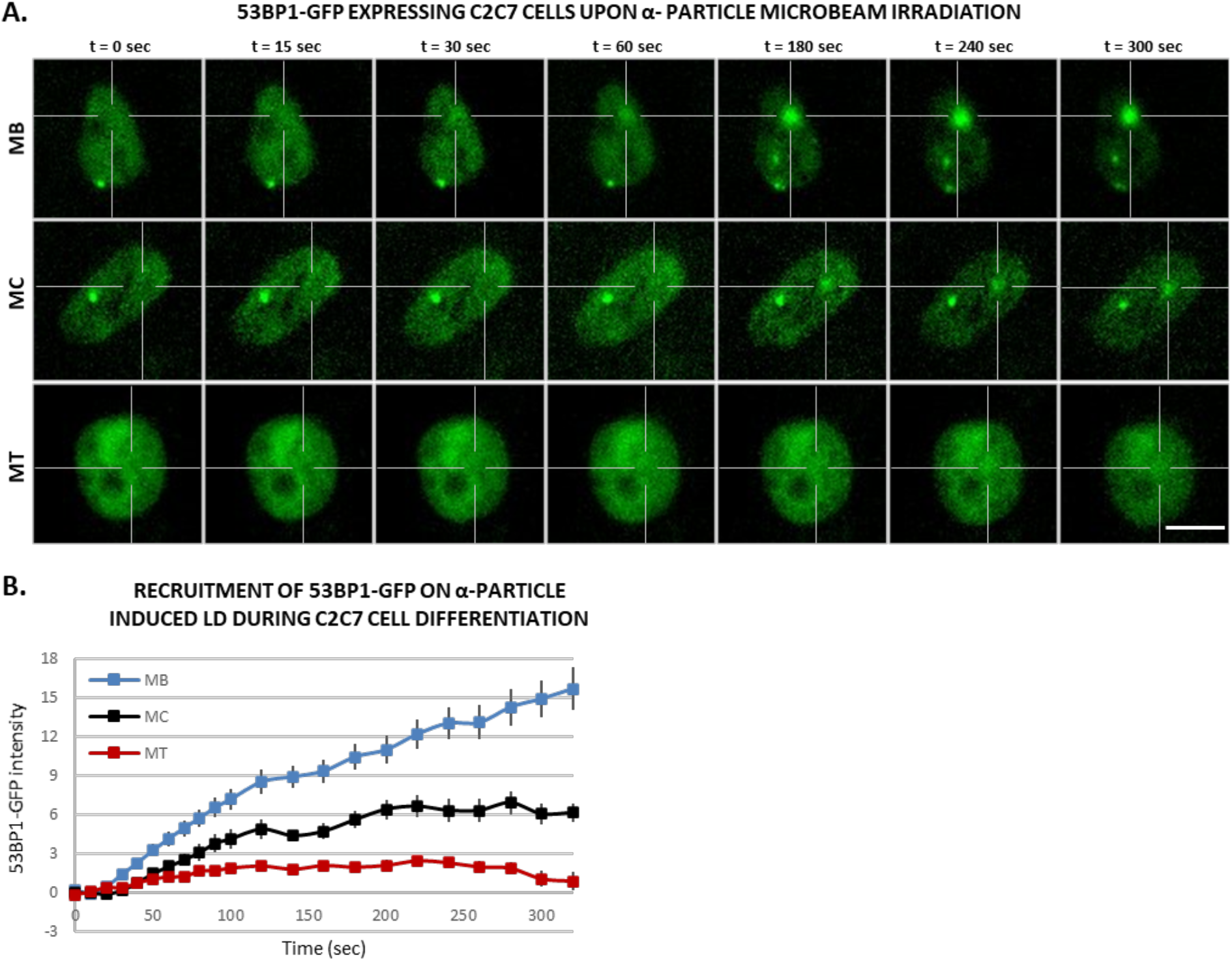
53BP1, a DDR factor, has almost no recruitment in multi-nucleated myotubes to local DNA damage. **A.** Sequential images of 53BP1-GFP recruitment to the locally damaged DNA (LD) by α-particle microbeam irradiation in stably 53BP1-GFP expressing C2C7 myoblasts (MB, upper panel), myocyte (MC, middle panel) and myotubes (MT, lower panel). The damaged areas are underlined by a dotted cross in the nucleus. Scale bar, 10 µm. **B.** Recruitment curve of 53BP1-GFP to the LD by α-particle microbeam irradiation in stably 53BP1-GFP expressing C2C7 MB (blue curve) in growth medium, MC (black curve) in differentiation medium (DM) and MT (red curve) in DM. The irradiation was applied at t = 10 s. N ≥ 3 independent experiments with 26-100 nuclei/cell type, mean ± SEM.

Taken together, our data obtained with local laser or α-particles irradiation pinpoint that the decrease of 53BP1 requirement is an intrinsic characteristic of differentiated myonuclei.

## DISCUSSION

Biochemical and genetic studies have provided valuable insights into the mechanism of action of DNA Repair and Transcription. However, despite almost three decades of structural, biochemical and cellular studies devoted to understand these fundamental cellular processes, many questions remained unanswered. One of these questions is how non replicative highly differentiated cells repair their genome, preserving their cellular functionality. Amongst highly differentiated cells, skeletal muscle fibers are a perfect example of post-mitotic cells and a model to study how DNA damage is repaired in cells that do not divide anymore. Exploring how DNA damage is repaired in skeletal muscle fibers is also highly relevant for human health as musculoskeletal injuries have been reported as late effects of treatment by radiation therapy ^54, 55^. These injuries include contractures, pain with motion, loss of muscle function and muscle weakness, requiring orthopedic appliances and reducing patients’ quality of life ^56–58^. For instance, in patients with lung cancer, breast cancer or stomach cancer, the chest wall and diaphragm are in the field of radiation treatment and in most occurrences, a decline in diaphragm efficiency is observed ^59^, negatively affecting quality of life.

We here studied how both oxidative damage and DSBs are repaired in myoblasts versus myonuclei in fibers, these two damages represent the most common DNA damage induced by radiotherapy. Additionally, oxidative damage is also endogenously induced by Reactive Oxygen Species (ROS) produced by the normal cellular metabolism and studying the oxidative damage repair capacity of skeletal muscle fibers might enlighten us on the progressive aging of this tissue.

Oxidative damage is repaired by the BER pathways that comes in two flavors: the short-patch (SP-BER) in which a single nucleotide gap is generated and subsequently filled and ligated and the long-patch repair (LP-BER) in which a gap of 2–10 nucleotides is generated and filled ^12^. We here used a previously produced knock-in mouse model that expresses endogenously a fluorescent tagged version of the protein FEN1 ^15^. FEN1 is a FLAP-endonuclease that plays a role in replication, processing the 5’ ends of Okazaki fragments in lagging strand DNA synthesis ^60^. In DNA repair FEN1 participates in the last steps of the LP-BER by removing 5’ overhanging flaps of DNA ^61^. For this reason, FEN1 is used as a *bona fide* marker of BER kinetics ^15^. Briefly, after damage induction, the accumulation of FEN1 correlates proportionally with the substrate produced by the BER reaction and the disappearance of the FEN1-YFP signal correlates with the disappearance of the substrate coinciding with the end of BER. We induced local oxidative damage (LOD), within nuclei of myoblasts and myotubes, with different damaging techniques that have been previously shown to produce different kind of DNA lesions ^38, 62^. We observed a clear reduction of the accumulation of FEN1 in LOD in myotubes versus myoblasts, and we could show that the turnover rate of FEN1 on LOD in myotubes is very slow compared to the one measured in myoblasts and previously measured in MEFs^15^. In these replicative cells, FEN1 proteins rapidly bind to and dissociate from the DNA flaps formed as intermediates in LP-BER. Different hypothesis can explain this result, we explore the possibility that the expression level of FEN1 and other BER factors might explain the reduced BER kinetics observed in fibers myonuclei. As we have previously shown in ^15^, FEN1 steady state levels are reduced in non-replicative cells (neurons, hepatocytes), we compared the concentration of FEN1 in myoblasts and myotubes and observed that myotubes have a lower FEN1 level, nevertheless we could show that this decrease does not statistically correlate with the reduction in the accumulation of FEN1 on LOD. It has been previously found that XRCC1, LIG1 and LIG3 have reduced expression levels in myotubes ^63^. We have verified and completed this study by quantifying FEN1, PARP1, APE1, XRCC1, LIG1 and found that all these proteins have lower steady state levels in myotubes versus myoblasts, arguing that LP-BER might be retarded in myotubes. While we clearly show that LP-BER is hindered in myotubes, we cannot exclude that SP-BER is indeed functional and might account for most of the repair reactions in these cells.

Beside oxidative damage, IR exposure induces DNA double strand breaks (DSBs) that are one of the most dangerous lesions for cells, because, if unrepaired or misrepaired, DSBs can lead to cell death or tumorigenesis ^39, 43, 44^. The canonical non-homologous end-joining (NHEJ) and homologous recombination (HR) are the two principal pathways to repair the majority of DSBs ^43, 44^. The HR pathway, requiring the presence of a homologous sequence on the sister chromatid to guide the repair, only occurs in late S and G2 phases, whereas NHEJ consisting in the re-joining of DSB ends, operates at all stages of the cell cycle ^45, 64^. Thus, we assessed whether the DSB repair mechanisms are delayed during myogenic differentiation as observed for the BER pathway. As the fusion of GFP to RAD51 affects HR-mediated DSB repair ^65^ we studied the recruitment of this central HR protein, which promotes the search for homology and strand pairing steps, by immunostaining. We observed no IR-induced foci (IRIF) of RAD51 in myotubes in contrast to myoblasts, and we did not detect RAD51 in myotubes by immunostaining, as previously reported by RT-qPCR and western-blot in ^22^. This finding is consistent with the fact that in myotubes the homologous donor sequence is not present. In addition, it also suggests that in post-mitotic cells, as myotubes, the sequence on homologous chromosome is rarely used. So far, our data lead us to hypothesize that the impairment of BER and HR pathways in myotubes is due to the decline of expression levels of some BER proteins and HR proteins, it remains an open if it could be related to the implication of these proteins in the replication-related processes.

Furthermore late IRIF formation of RAD51 in mono-nuclear cells (i.e., myoblasts and myocytes) suggests that NHEJ is the major repair pathway in the initial response to irradiation-induced DNA lesions and that HR takes place later, in our present study 2 hours post-Irr, in accordance with the literature ^67^. The absence of RAD51 in post-mitotic myotubes confirms that the DSB repair occurring in myotubes carried out in an NHEJ-dependent manner, as was expected but not formally proven yet. To further understand the DSB repair capacity of proliferating myoblasts ^68^ and post-mitotic myotubes ^69^, we concentrated on NHEJ factors upon induced local DNA damage. NHEJ ensures the direct ligation of broken ends without the need for a homologous template and operates both in non-dividing cells and proliferative cells. Here we used the myogenic C2C7 cell line, stably expressing GFP-tagged NHEJ proteins, KU80 and DNA Ligase 4. KU80 with the KU70 protein forms the heterodimer KU that initiates NHEJ by recognizing and binding DNA ends and subsequently by recruiting the catalytic subunit of the DNA-dependent kinase (DNA-PKcs), leading to the formation of the DNA-PK holoenzyme ^19^. Recruited by DNA-PK, LIG4 with its co-factors XRCC4 and XLF/Cernunnos acts at the later step of NHEJ to perform the ligation of processed DNA ends ^19^. In the present study, we used these two proteins as markers of NHEJ kinetics, covering initial and final steps of NHEJ-mediated DSB repair. Our data suggest that NHEJ-dependent DSB repair machinery is also weakened in myotubes in agreement with observed reduced recruitment of NHEJ factors, KU80 and DNA Ligase 4, to induced local DNA damage upon α-particle or laser-irradiation in comparison to mono-nuclear myogenic cells. In contrast to the BER and HR mechanisms, we could not link this reduction in NHEJ activity to a reduced steady state level of NHEJ proteins, since myotubes present at least for the KU complex and LIG4 a higher expression level. However, we observed that the turnover rate of KU80 on local DNA damage sites induced by laser-irradiation is faster in myotubes compared to the one measured in myoblasts, suggesting a reduced occupancy onto the damaged substrate that could explain the decreased KU80 and LIG4 kinetics observed in myotubes.

Upon DNA damage, the H2A histone variant, H2AX gets phosphorylated at serine 139, then called γ-H2AX ^70^, and acts as signaling machinery to induce chromatin relaxation and as a scaffold for the DNA repair factors at the proximity of DNA damage. Shortly after 53BP1 is recruited to DNA lesions, forming IRIF and favors NHEJ by its inhibitory effect on broken DNA end resection induced by MRN complex as well as its role in heterochromatin relaxation. Upon DNA damage the appearance and disappearance of γ-H2AX and 53BP1 IRIF along with repair of DNA lesions, makes them good DSB markers to assess repair kinetics ^41, 42^. Thereby assessing γ-H2AX and 53BP1 IRIF upon X-ray irradiation, we showed that DSB repair kinetics are declined in myotubes. Despite the same doses of irradiation, we have observed that initial γ-H2AX foci number were higher in proliferating myoblasts than post-mitotic myotubes, which could most likely be due to DNA copy number as asynchronously proliferating myoblasts have cells in S and G2 phases with replicated DNA, in agreement with previously reported paradigm ^71^. Nevertheless, the disappearance of DSB markers in MB was higher over time and the IRIF reached lower number in comparison to MT 24 hrs post-Irr, suggesting a faster repair kinetics in MB. Indeed, the notion of DNA copy number-dependent DDR could be another reason that myoblasts have a greater recruitment capacity of NHEJ factors, such as KU80 and LIG4, in comparison to myotubes, which have nuclei only in G0 state with 1 copy of DNA.

Moreover, ionizing radiation can induce DSBs in direct and indirect manners through cumulated SSBs at close proximity and excitation by oxidative radicals produced upon radiation. High LET radiation as α-particles is reported to produce more complex DSBs as well as clusters of DNA lesions in comparison to low LET X-ray radiation-induced DSBs, which are induced in a more dispersed manner ^36^. Thus, taking into account the induced SSBs, base lesions and clustered DNA damages upon radiation in MT along with reduced DNA SSB repair machinery could be one reason for reduced DSB repair through NHEJ. As the proliferating MBs have all the DNA damage repair machineries available, they could process the SSBs, and other damages followed by DSB repair whereas MTs could be stalled or slowed at the initial process of SSB repairs before DSB repair. Future works, complementary to this study, will be necessary to identify the factor inducing a decrease in DSB repair through NHEJ in MT.

Additionally, 53BP1, one of the initial players in DSB repair, is clearly recruited to the local DNA damage site upon α-particle or laser-irradiation in myoblasts whereas in myotubes, we observed very low recruitment of 53BP1, in agreement with our findings in myogenic cells upon X-ray irradiation. One of the roles for 53BP1 upon IR-induced DNA damage is to inhibit MRN complex initiated DNA-end resection and favor NHEJ in competition with HR factor BRCA1 ^30, 52^. Accordingly, upon myogenic differentiation cells exit cell cycle and generate post-mitotic myotubes ^72^ and additionally the absence of HR, predicted from the absence of RAD51 in myotubes, suggests that 53BP1 is not an essential protein for the initial DNA damage response in myotubes. Besides it has been reported that upon DNA damage, 53BP1 modulates p53-dependent and -responsive genes, for instance cell cycle and pro-apoptotic targets ^73^ although myotubes have no activation of p53 upon Irr-induced DNA damage ^66^, which could also explain strongly reduced 53BP1 response to induced DNA damage in myotubes. An additional and essential role of 53BP1 is during the repair of DSBs at heterochromatin structures, which is reported as slow kinetic repair. 53BP1 is necessary for ATM localization at the damage site and phosphorylation of KAP1 leading for chromatin relaxation ^46^. In addition, previously it was suggested that 53BP1 have a role of protecting the DNA broken ends independent to ATM, thus from translocations ^74^. Consequently, the role of 53BP1 in chromatin relaxation at latter slow kinetic DSB repair and protection of broken DNA ends could explain the late recruitment of 53BP1 to the DNA damage site observed in MT.

For the first time, we systematically analyzed major DNA repair mechanisms of IR-induced lesions, BER, HR and NHEJ, along myogenic differentiation. We found that in the most differentiated myogenic cells, myotubes, all of these mechanisms present weakened kinetics of recruitment of DNA repair proteins at IR-damaged DNA. For BER and HR, this decline can be link to a reduced need for these proteins since myotubes no longer replicate their DNA. However, the factor responsible for this decline in NHEJ has yet to be identified.

## MATERIALS AND METHODS

### Primary cell isolation and myogenic cell culture

Muscle stem cells were freshly isolated from the hind limbs of neonatal (4-6 days old) C57B/6J mice or *Fen1-YFP* mouse strain ^15^ as previously described ^75^. Briefly, the hind limb muscles were chopped off and digested by a mix of 4.8 U/mL Dispase II (neutral Protease, grade II) and 0.4 % Collagenase A in DMEM Glutamax. After a pre-plating step followed by a centrifugation at 600 G for 10 min, the cell pellet was resuspended in myogenic cell medium (DMEM/F12 1:1 (GIBCO), 20 % FBS (EUROMEDEX), 1 % Penicillin/Streptomycin (GIBCO), 0.5 % Gentamicin (GIBCO) and 2 % Ultroser G (PALL)) and immediately seeded on cell dishes pre-coated with 0.1 mg/ml of Poly-D-Lysine (SIGMA) and Matrigel (Corning). The day after the entire medium was refreshed, then 50 % of the cell medium was refreshed the 3rd day post-seeding and every consecutive day after for inducing the myogenic differentiation and generating myotubes in culture.

The immortalized myogenic C2C7 cells ^40^ were cultured in the similar conditions as primary myogenic cells in growth medium (GM) containing 20 % FBS, and 1 % P/S in DMEM Glutamax (GIBCO) and upon reaching ≥ 80 % confluency, the medium was switched to differentiation medium (DM) containing 2 % Horse Serum (HS, GIBCO), and 1 % P/S in DMEM Glutamax, 50 % of DM was refreshed after 3 days and every consecutive day after.

All the cells were incubated in a humidified atmosphere at 37 °C with 5 % CO2 and 3 % O2.

### Fibroblast culture and PEG fusion

Primary FEN1-YFP fibroblasts were isolated as previously described from *Fen1-YFP* mice ^15^ and incubated in 15 % FBS, 1 % P/S in DMEM Glutamax at 37 °C with 5 % CO2 and 3 % O2 in a humidified atmosphere. When indicated, fibroblasts were fused by incubating the cells in 50% (vol/vol) PEG4000/DMEM Glutamax for 10 min at 37 °C followed by further incubation of cells minimum for 24 hrs in normal culture conditions.

### Plasmids and transfections

The plasmids expressing GFP-tagged protein of interest were kindly provided by Pascale Bertrand (53BP1-GFP; CEA, iRCM/IBFJ, UMRE008 Stabilité Génétique, Cellules Souches et Radiations, Fontenay-aux-Roses, France), Dik C. van Gent (KU80-GFP; Departments of Cell Biology and Genetics, Erasmus MC, Rotterdam, The Netherlands) and Mauro Modesti (Ligase4-GFP; CRCM, CNRS UMR7258, Inserm U1068, Institut Paoli-Calmettes, Aix-Marseille Université, Marseille, France). These plasmids were transfected on both primary and C2C7 myoblasts with TurboFect (ThermoFisher Scientific) according to the manufacturer’s instructions. In order to have successful transfection, primary cells were transfected 3 days post-seeding, which provided enough time for quiescent satellite cells to activate and expand in culture, whereas C2C7 cells were transfected 1 day post-seeding at about 50 % - 60 % confluency. Then, subsequent experiments were performed 24 hours – 1week post-transfection on primary cells.

For C2C7 cells, 24 hours post-transfection the GFP-expressing cells were enriched under geneticin (G418 sulfate) (GIBCO) selection for 10 consecutive days. Then, the GFP-tagged protein expressing C2C7 cells were isolated by FACS, which provided us stable and homogenous GFP-tagged protein expressing C2C7 lines, which were further expanded.

### DNA damage induction

#### X-Ray

***X-*** Cells were irradiated with 5 Gy of X-ray (medical linear accelerator, Elekta Synergy Platform, Elekta SAS, Boulogne-Billancourt, France; 10 MV; dose rate 3 Gy.min^−1^ in air kerma free in air) in the X-ray irradiation platform of IRSN, Fontenay-au-Roses.

#### MIRCOM, Microbeam α-particle radiation

We performed the irradiation of samples with α-particles by using the MIRCOM facility, operated by the Institute for Radiological Protection and Nuclear Safety (IRSN) in Cadarache, France ^37^. The low energy charged α-particles have low penetration capacity and thus short travel distance through matter, therefore the cells are seeded in a special cell dish with a 4 µm thick polypropylene foil (Goodfellow) ^76^. To provide optimal cell growth conditions, the polypropylene foil is pre-coated with 10 ng/µl of Cell-Tak (Corning, Fisher Scientific) followed by Matrigel (Corning, Fisher Scientific). Then cells are placed under an inverted epifluorescence microscope (AxioObserver™ Z1, Carl Zeiss Microscopy GmbH, Jena, Germany) within a 37 °C heating chamber. The nuclei of the cells are identified and selected for irradiation with a 20X objective (Zeiss LD Plan-NEOFLUAR 20x/0.4 Korr). The microbeam extraction occurs on the other side of the cell dish where the microbeam of α-particles is aligned and focused by 4 magnetic quadrupoles in a “Russian quadruplet” configuration. Microbeam of 6 MeV α-particles is generated by a 2 MV Tandetron™ accelerator manufactured by High Voltage Engineering Europa B.V. (HVEE, Amersfoort, The Netherlands), then sent on the targeted zone by electrostatic scanning plates in desired number of ions or for desired amount of time, as previously described ^37, 38^. To follow the recruitment kinetics of GFP-tagged proteins, we started time-lapse imaging 10s before irradiation and recorded images every 2 s with a monochromatic AxioCam™ MRm rev. 3 CCD camera (Carl Zeiss Microscopy GmbH, Germany) using the CRionScan software. We recorded images with an exposure time of 800 ms. In total, we kept cells in the microbeam chamber for less than 30 min.

#### Local damage induction with multi-photon laser

Cells were seeded onto coverslip. Imaging and local damage induction were performed on confocal Zeiss 980 (CRCL, Lyon) coupled with a bi-photon 800nm laser confocal LSM780NLO Zeiss microscope (IRSN, Fontenay-aux-Roses) coupled with a bi-photon 800 nm laser (Chameleon Vision II, Coherent). The local DNA damage was obtained with 800 nm pulsed output at 10 % power. To target cells, 30 pixels circular regions (or 10 x 1 µm rectangular form, when indicated) is used to induce DNA damage in nucleus with 13 ms of exposure.

### FRAP on local DNA damage induced by multiphoton laser

Imaging and FRAP were performed on Confocal Zeiss 980 (CRCL, Lyon). 488 nm laser at 100 % intensity and 1 iteration is used to induce photo-bleach on multiphoton laser damage. The bleach is realized after that maximum fluorescence intensity of LD is achieved.

### Immunofluorescence labelling and image analysis

Upon DNA damage induction, the samples were fixed with 2 % paraformaldehyde (PFA, EMS, Euromedex) in phosphate-buffered saline (PBS, Gibco) for 20 min respective to indicated time-points, followed by permeabilization with 0.5% Triton X-100 in PBS for 5 mins. To increase the stringency the samples were washed with 0.1% Tween 20 in PBS for 20 min, then blocked with 5% BSA (Sigma-Aldrich), 0.1% Tween 20 in PBS. The samples were incubated with the indicated primary antibodies overnight at 4°C and with the appropriate fluorophore-conjugated secondary antibodies for 1 hr at room temperature (RT). Finally, the samples were incubated with DAPI (1/25,000 in PBS) for 5 min and mounted with Prolong Diamond antifade mounting medium (Invitrogen). The samples were imaged and analyzed with C-Plan Apochromat 63x/1.4 Oil DIC M27 objective under confocal microscope (LSM780NLO, Zeiss).

### Antibodies

Primary antibodies used during immunofluorescence (IF) experiments are: rabbit anti-53BP1 (Bio-Techne, Novus Biologicals, NB 100-304; 1:500); mouse anti-APE1(Abcam, clone 13B8E5C2, ab 194; 1:500); rabbit anti-FEN1(Abcam, ab 17993; 1:500); mouse anti-XRCC1(Abcam, ab 1838; 1:50); mouse anti-γH2AX (Millipore, UpState, 05-636; 1:2,000); rabbit anti-KU70/80 (Abcam, ab 53126; 1:400); mouse anti-DNA Ligase 1 (Sigma Aldrich, Merck, clone 5H5, MABE1905; 1:500); mouse anti-PARP1 (R&D Systems, Bio-Techne, 4338-MC; 1:1,000); rabbit anti-RAD51 (Abcam, ab137323; 1:400).

Secondary antibodies used are: donkey anti-mouse or donkey anti-rabbit coupled to Alexa Fluor 488, 594 or 647 (Invitrogen, ThermoFisher; 1:1,000), anti-mouse coupled to Alexa fluor 594 (Invitrogen, A-11005) and anti-rabbit coupled to Alexa fluor 488 (Invitrogen, A-11008).

### Quantification and Statistical analysis

All the images were processed, analyzed, and quantified by software ImageJ (version 1.53e) (7) and statistical analyses were performed by software Prism version 9 (GraphPad Inc.) and Excel (Microsoft). In order to quantify the fluorescence re-localization of GFP-tagged proteins observed with time-lapse imaging upon multiphoton laser damage or MIRCOM irradiation, we manually selected and followed regions of interest (ROI). We measured the mean intensity of ROIs in every image and plotted them against time. Then obtained data were corrected for non-specific fluorescence bleaching and normalized for the fluorescence intensity measured before irradiation. For FRAP on local DNA damage induced by multiphoton laser, mobility curve show relative fluorescence (fluorescence post-bleach divided by fluorescence pre-bleach) plotted against time. All statistical analyses were performed from at least 2 independent experiments.

## ACKNOWLEDGEMENTS

The authors thank the PSE-SANTE/SDOS/LMDN team of the IRSN for their excellent technical expertise on the MIRCOM facility; Valérie Buard (PSE-SANTE/SERAMED/LRMEd) for her technical support on the IRSN confocal microscopy facility, Yoann Ristic and Miray Razanajatovo (PSE-SANTE/SDOS/LDRI) for dosimetry and X-ray irradiation; Lyon microscopy facility (CIQLE, Lyon) and mouse house (ALECS, Lyon). We are also grateful to Dr. Gaetan Gruel and Dr. Stéphane Illiano for their critical reading of the manuscript.

## SUPPLEMENTAL MATERIALS AND METHODS

### Inhibitors

When indicated, DNA repair inhibitors were added in cell culture medium 3 hrs before DNA damage induction and experiments were performed in the presence of inhibitors. VE821 (Sigma-Aldrich, SML1415) and KU55993 (Sigma-Aldrich, SML1109) were used at 5 µM and stock solutions were 1 mM diluted in DMSO. All the cells were incubated in a humidified atmosphere at 37 °C with 5 % CO2 and 3 % O2.

### NIH-3T3 cell culture and PEG fusion

NIH-3T3 cells (CRL-1658) were obtained from the American Type Culture Collection (ATCC, LGC Standards) and maintained in 10% iron fortified calf bovine serum (ATCC, LGC Standards), 1% P/S in DMEM Glutamax in a humidified atmosphere at 37 °C with 5 % CO2 and 20 % O2. When indicated, NIH-3T3 cells were fused by incubating the cells in 50% (vol/vol) PEG4000/DMEM Glutamax for 10 min at 37 °C followed by further incubation of cells minimum for 24 hrs in normal culture conditions.

### RRS assay

FEN1-YFP cells were grown and differentiate on fluorodish. RNA detection was done using a Click-iT RNA Alexa Fluor Imaging kit (Invitrogen), according to the manufacturer’s instructions. Briefly, cells were incubated for 2 hrs with 100 µM of 5-ethynyl uridine (EU). After fixation with 4 % paraformaldehyde (PFA) for 15 min at 37 °C and permeabilization with phosphate-buffered saline (PBS) and 0.5 % Triton X-100 for 20 min, cells were incubated for 30 min with the Click-iT reaction cocktail containing Alexa Fluor 594. After washing, cells are incubated with DAPI for 15 min. The samples were imaged with Zeiss Z1 imager right using a ×40/0.75 dry objective. The acquisition software is Metavue Using ImageJ, the average fluorescence intensity per nucleus was estimated after background subtraction and normalized to myoblasts.

### RNA fluorescence in situ hybridization (RNA FISH)

FEN1-YFP cells were grown on fluorodish, washed with warm PBS, and fixed with 4 % PFA for 15 min at 37 °C. After two washes with PBS, cells were permeabilized with PBS + 0.4 % Triton X-100 for 7 min at 4 °C. Cells were washed rapidly with PBS before incubation (at least 30 min) with prehybridization buffer: 15 % formamide in 2× SSPE (Sodium Chloride-Sodium Phosphate-EDTA) (0.3 M NaCl, 15.7 mM NaH2 PO4 ·H2O, and 2.5 mM EDAT [Ethylenediaminetetraacetic acid] et pH8.0). 35 ng of the probe was diluted in 70 μl of hybridization mix (2× SSPE, 15 % formamide, 10 % dextran sulfate and 0.5 mg/mL tRNA). Hybridization of the probe was conducted overnight at 37 °C in a humidified environment. Subsequently, cells were washed twice for 20 min with prehybridization buffer and once for 20 min with 1× SSPE. After extensive washing with PBS, the coverslips were mounted with Vectashield containing DAPI (Vector). The probe sequence (5′ to 3′) is Cy5-AGACGAGAACGCCTGACACGCACGGCAC. The samples were imaged with Zeiss Z1 imager right using a ×40/0.75 dry objective. The acquisition software is Metavue Using ImageJ, the average fluorescence intensity per nucleus was estimated after background subtraction and normalized to myoblasts.

### Immunofluorescence protein level quantification

For BER proteins, image acquisition has been performed on a Zeiss Z1 imager right using a ×40/0.75 dry objective. The acquisition software is Metavue. Using ImageJ, the average fluorescence intensity per nucleus was estimated after background subtraction.

For DNA double strand break (DSB) repair proteins, endogenous protein content in myogenic cell sub-populations was quantified by subtracting the mean fluorescent intensity of background from mean fluorescent intensity of total nuclei detected by the indicated antibodies or of Ligase 4-GFP from the images of non-irradiated cells acquired by confocal LSM780NLO Zeiss microscope (IRSN, Fontenay-aux-Roses).

### TUNEL assay

To label DSB and apoptosis upon irradiation, we performed Click-iT plus TUNEL assay (Invitrogen, ThermoFisher) following the manufacturer’s protocol. After 2 % PFA fixation and 0.5 %Triton X-100 permeabilization of cells, samples were treated with TdT (terminal deoxynucleotidyl transferase) enzyme and label 3 ′ OH of the DNA broken ends with EdUTP, followed by Click-iT UTP labelling with fluorophore for fluorescent detection.

### **X-** ray irradiation settings

The cells were irradiated in 2 ml of cell medium in 12 well plates placed at the center of 30 cm x 30 cm of irradiation field. Cells were irradiated with 5 Gy of X-ray (medical linear accelerator, Elekta Synergy Platform, Elekta SAS, Boulogne-Billancourt, France; 10 MV; dose rate 3 Gy.min^−1^ in air kerma free in air, distance of the source :1.1 m) in the X-ray irradiation platform of IRSN, Fontenay-au-Roses. Uncertainty on the dose was estimated to be 5 %.

**Figure S1.**
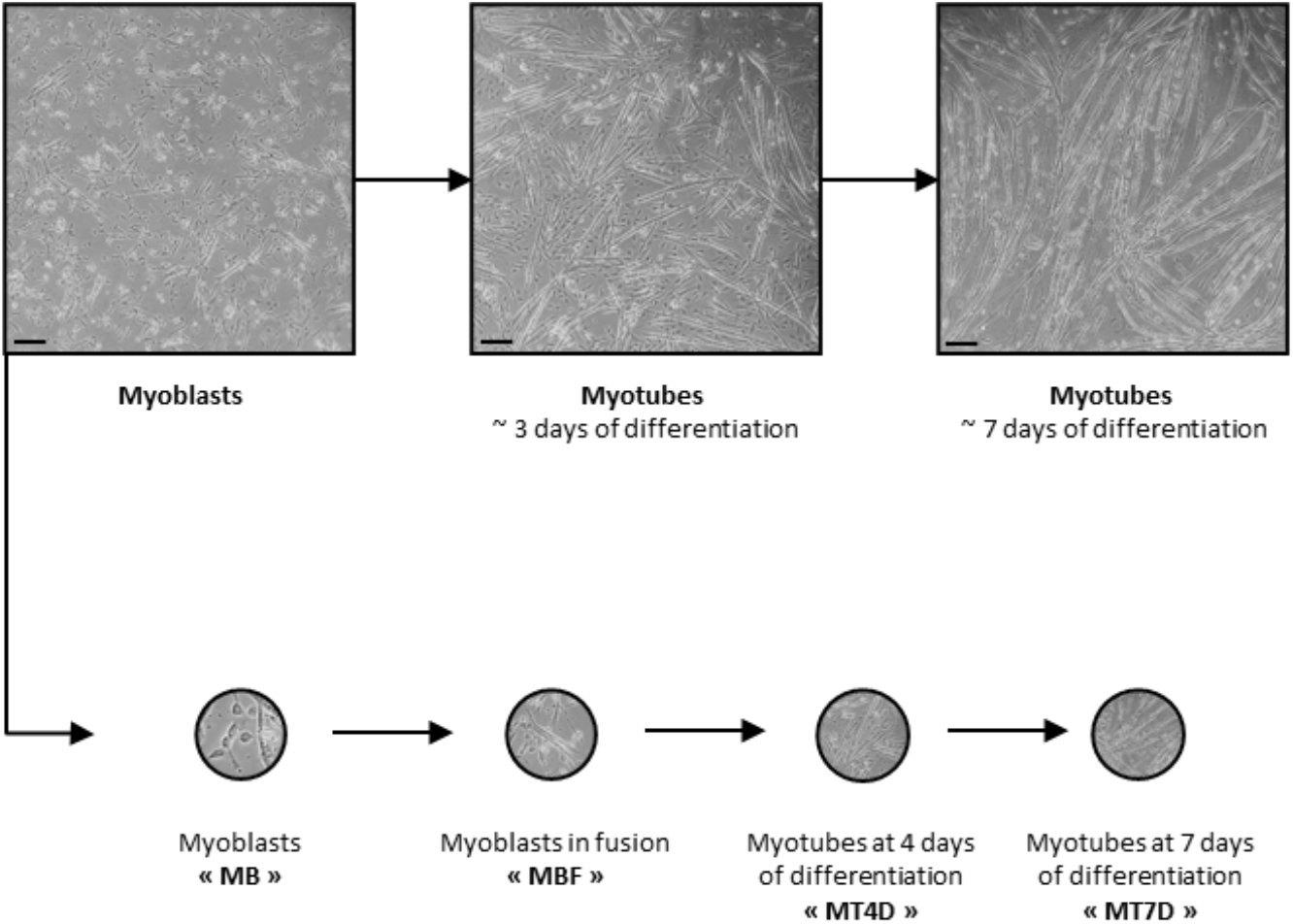
Four steps of FEN1-YFP myofibrillogenesis. Myoblasts (MB) are isolated from muscles of Fen1-YFP 5 days old mice. After 4-5 days in culture, proliferative myoblasts were induced to fuse (MBF) and differentiate into mature myotubes (MT). The first experiments were conducted with MB, MBF and two steps of differentiation: myotubes of 4 days (MT4D) and myotubes of 7 days (MT7D). Scale bar, 100 µm.

**Figure S2.**
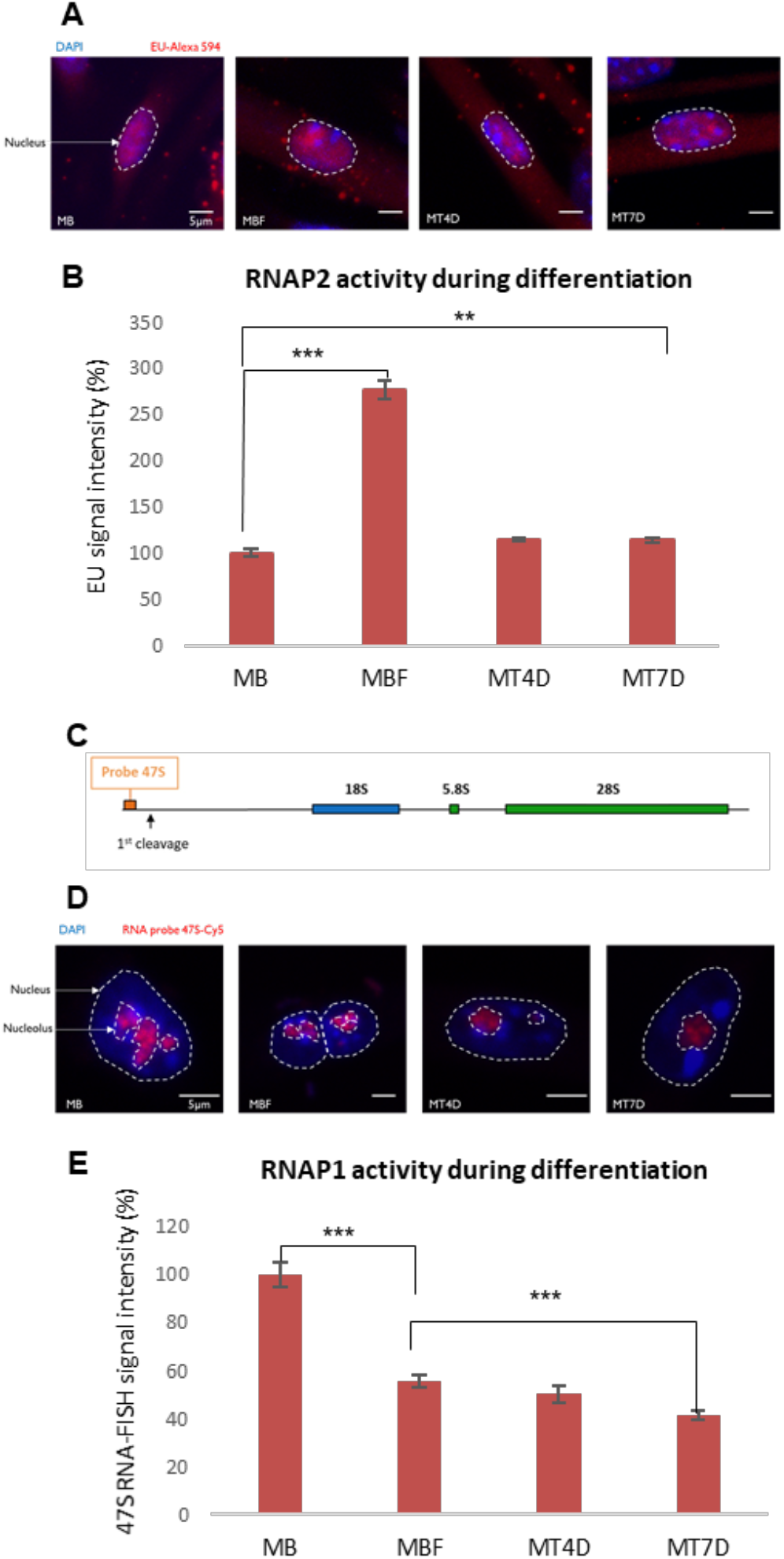
Transcriptional activity during myofibrillogenesis. **A.** Images of <u>5-ethynyl uridine (</u>EU) incorporation measuring mRNA production for each condition during differentiation, in myoblasts (MB) that were isolated from muscles of 5 days old mice, myoblasts that were induced to fuse (MBF), myotubes of 4 days (MT4D) and myotubes of 7 days (MT7D). DAPI signal is depicted in blue and EU incorporation is illustrated in red. Nuclei are delimited by dashed lines. Scale bar, 5µm. **B.** Quantification of RNA synthesis determined by EU incorporation in the course of myogenic differentiation, in MB, MBF, MT4D and MT7D. Error bars represent the SEM obtained from at least 50 cells and data are representative of three independent experiments, significance by t-Test. * p≤0.05, ** p≤0.01, *** p≤0.001. **C.** Schematic representation of rDNA unit and localization of the 47S pre-rRNA probe. **D.** Images of 47S RNA-FISH for each condition during differentiation, in MB, MBF, MT4D and MT7D. DAPI signal in blue and 47S probe is illustrated in red. Nuclei and nucleoli are delimited by dashed and dotted lines, respectively. Scale bar, 5 µm. **E.** Quantification of 47S Synthesis determined by 47S probe hybridization in the course of myogenic differentiation, in MB, MBF, MT4D and MT7D. Error bars represent the SEM obtained from at least 50 cells and data are representative of three independent experiments, significance by t-Test. * p≤0.05, ** p≤0.01, *** p≤0.001.

**Figure S3.**
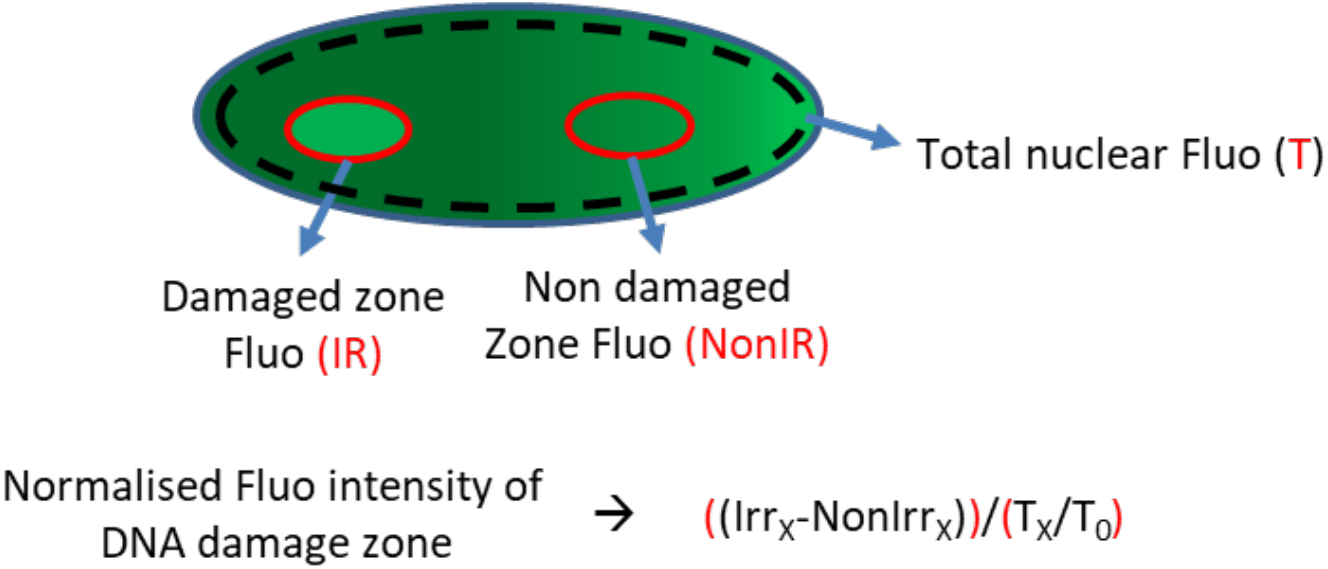
**Schematic representation** of zones selected for quantification of fluorescent protein-tagged protein intensity accumulated at the DNA damage site and the equation used for calculating the recruitment kinetics.

**Figure S4.**
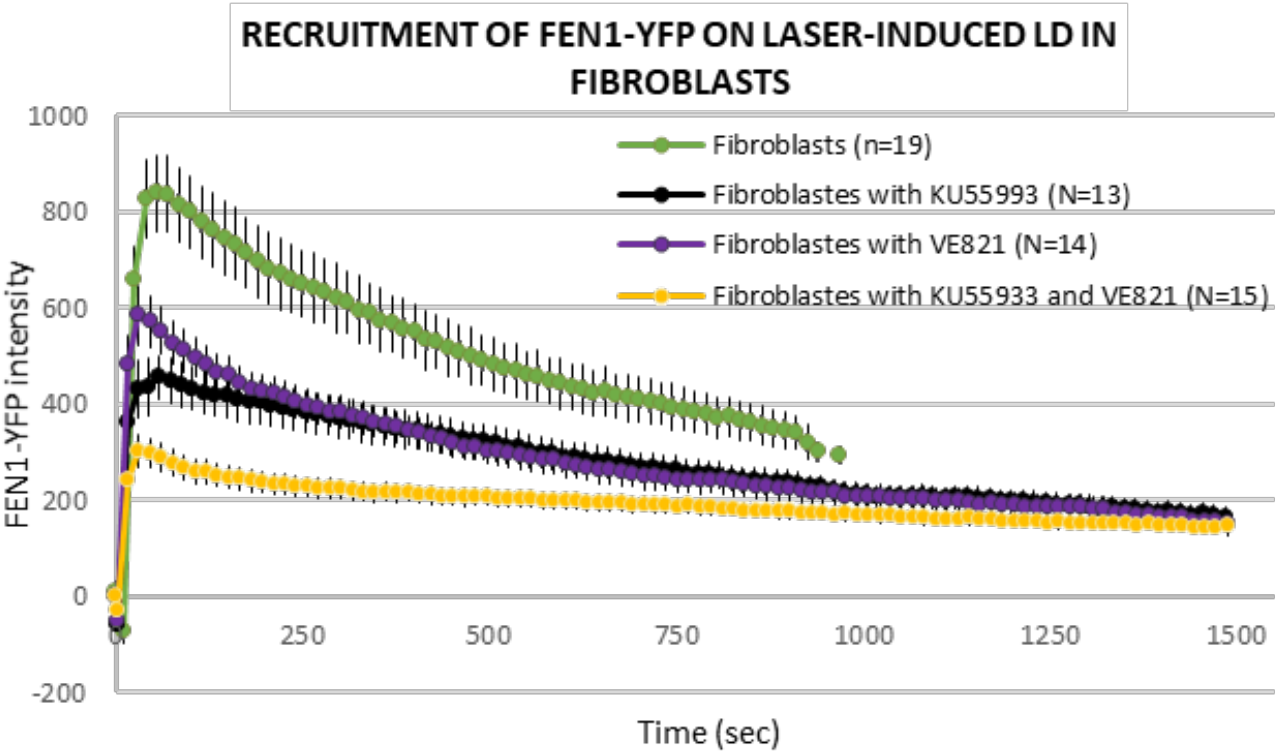
Dynamic behavior of FEN1-YFP upon local DNA damage induction by laser-irradiation after inhibition of DNA damage signaling. Recruitment curve of FEN1-YFP onto the locally damaged DNA (LD) by laser-irradiation in untreated primary dermal fibroblast isolated from the Fen1-YFP mouse model (green curve), treated with the ATM inhibitor Ku55993 (black curve), treated with the ATM/ATR inhibitor VE821 (violet curve) and treated with both Ku55993 and VE821 (yellow curve).

**Figure S5.**
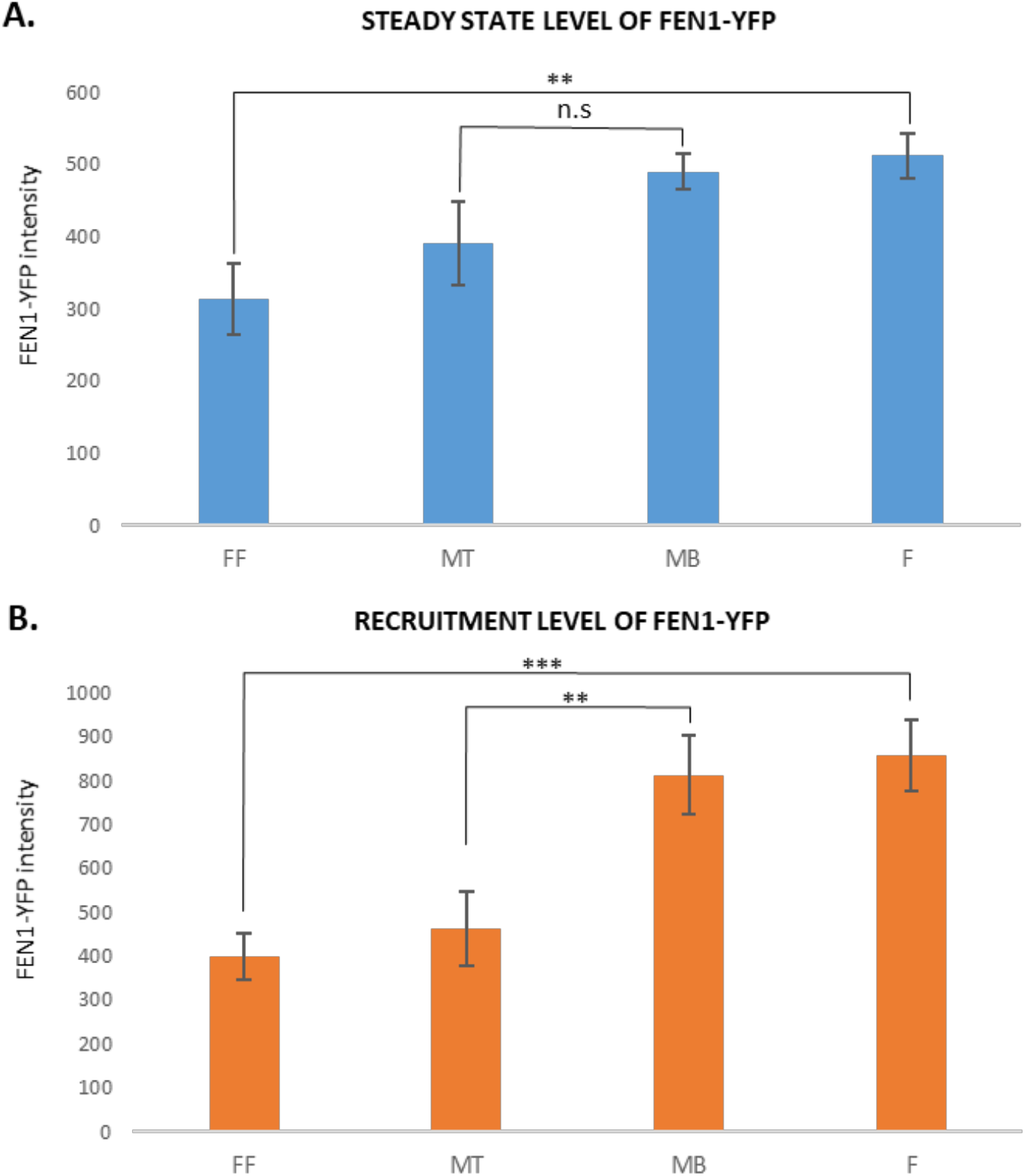
The reduction of DNA repair activity in differentiated FEN1-YFP myonuclei. **A.** The steady state level of FEN1-YFP represents the mean of YFP intensity before the locally damaging DNA (LD) by laser-irradiation in different cellular types isolated from the Fen1-YFP mouse model: myoblasts (MB) and myotubes (MT); primary dermal fibroblast isolated (F) and the same fibroblasts fused with PEG (FF). Error bars represent the SEM obtained from at least 15 nuclei, significance by t-Test. * p≤0.05, ** p≤0.01, *** p≤0.001. **B.** The level of FEN1-YFP recruited to the LD by laser-irradiation represents the mean of YFP maximum intensity after DNA damage induction in different cellular types isolated from the Fen1-YFP mouse model: MB and MT; F and FF. Error bars represent the SEM obtained from at least 15 nuclei, significance by t-Test. * p≤0.05, ** p≤0.01, *** p≤0.001.

**Figure S6.**
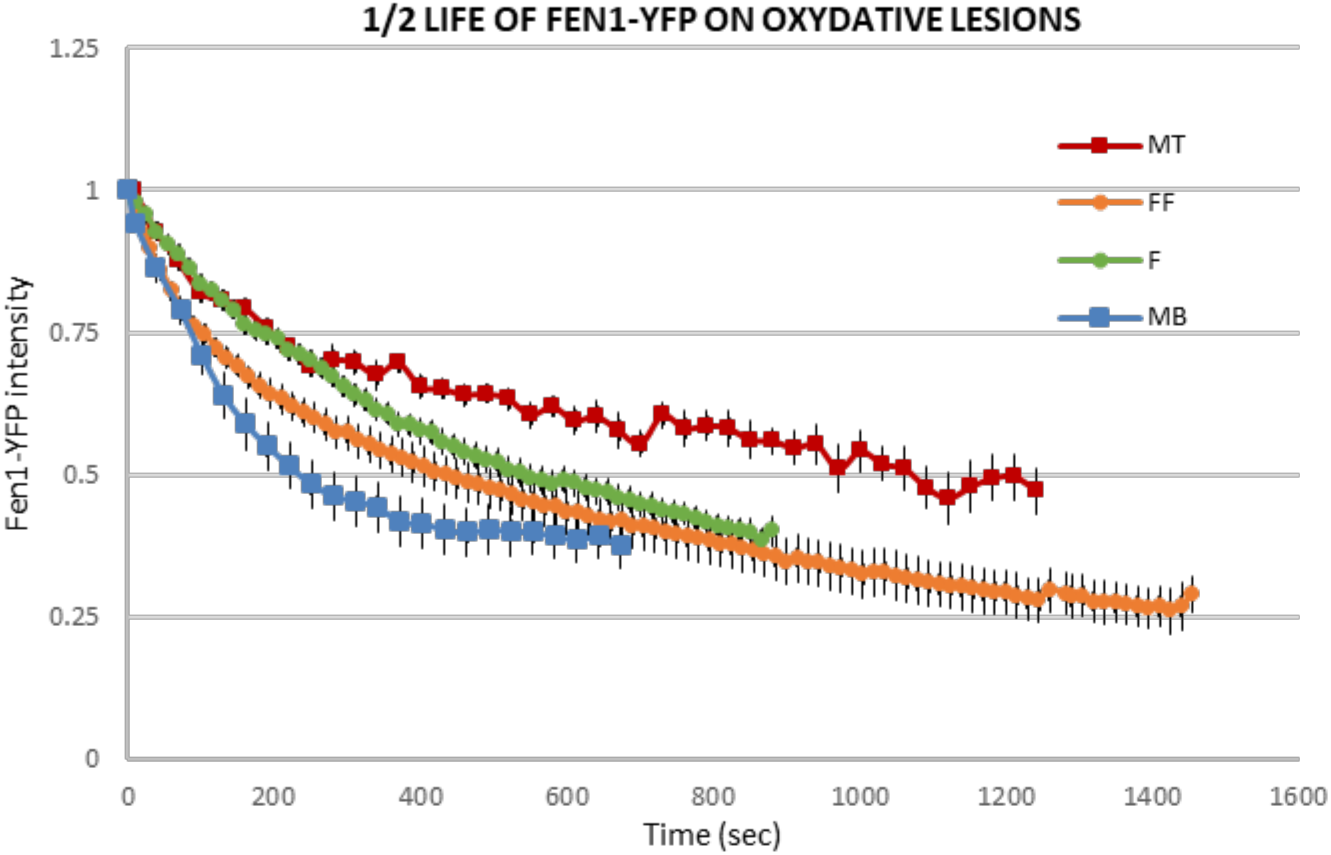
The reduction of DNA repair activity in differentiated FEN1-YFP myonuclei. Half-life curves of FEN1-YFP on oxidative lesions at the locally damaged DNA (LD) by laser-irradiation in different cellular types isolated from the Fen1-YFP mouse model: myoblasts (MB, orange curve) and myotubes (MT, red curve), primary dermal fibroblast isolated (F, green curve) and the same fibroblasts fused with PEG (FF, blue curve).

**Figure S7.**
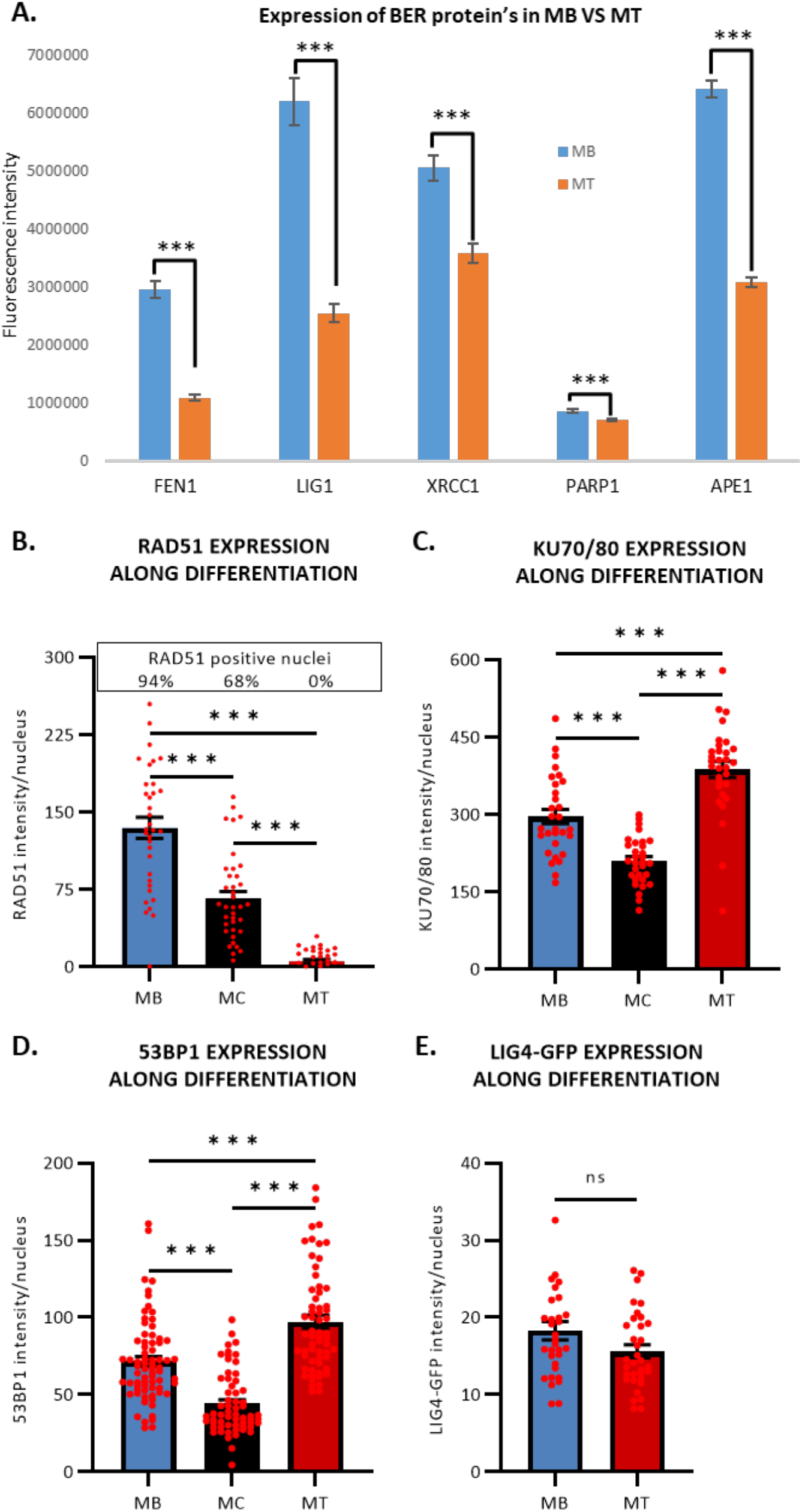
The reduction of BER and DNA Double Strand Break repair (DSBR) protein’s expression in myotubes. **A.** Quantification of BER protein expression level by immunofluorescence (IF) during myogenic differentiation, in myoblasts isolated from muscles of 5 days old mice and their committed progeny, myotubes (MT). We measured thank to the use of specific antibody the fluorescence intensity of FEN1, DNA Ligase 1 (LIG1), XRCC1, PARP1 and APE1 in MB (blue) and MT (red). Error bars represent the SEM obtained from at least 50 nuclei and data are representative of three independent experiments, significance by t-Test. * p≤0.05, ** p≤0.01, *** p≤0.001. **B, C, D.** Quantification of DSBR protein expression level by IF during myogenic differentiation, in C2C7 MB, myocytes (MC) and MT. We measured thank to the use of specific antibody the fluorescence intensity of RAD51, KU70/80 complex and 53BP1 in MB (blue), MC (black) and MT (red). Error bars represent the SEM obtained from at least 30 nuclei/cell type and data are representative of three independent experiments. Significance by 1-way ANOVA with post-hoc Tukey’s multiple comparison test, * p≤0.05, ** p≤0.01, *** p≤0.001. **E.** Quantification of LIG4-GFP protein expression level by IF during myogenic differentiation, in C2C7 MB and MT. Due to the absence of commercial antibody against murine DNA ligase 4 (LIG4) working in IF, we measured the GFP fluorescence intensity of LIG4-GFP in MB (blue) and MT (red). Error bars represent the SEM obtained from at least 30-35 nuclei/cell type and data are representative of three independent experiments. Significance by unpaired t-Test, * p≤0.05, ** p≤0.01, *** p≤0.001.

**Figure S8.**
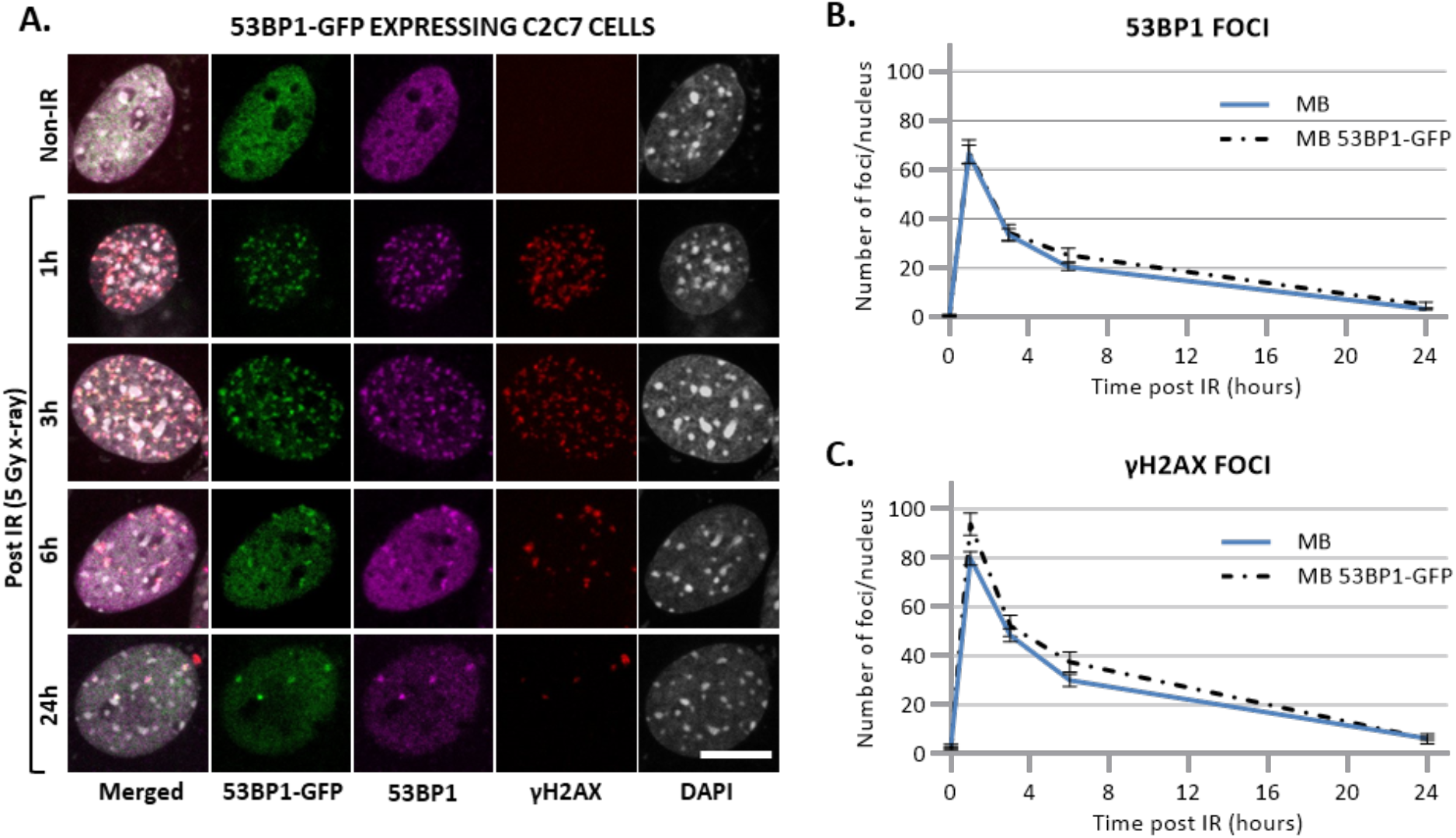
Disappearance of DSB markers upon X-ray Irradiation. **A.** Representative images of stably 53BP1-GFP expressing C2C7 myoblasts at the indicated time after 5 Gy of x-ray irradiation immunolabelled with antibodies against DSB markers: 53BP1 (magenta) γH2AX (red). DNA was stained with DAPI (grey). Scale bars, 10 µm **B, C.** Quantification of 53BP1 foci (B) and γH2AX foci (C) per nucleus in C2C7 myoblasts, un-transfected (blue curve) or stably transfected with 53BP1-GFP expressing plasmid (dashed black curve) at the indicated time upon 5 Gy of X-ray irradiation. N=1 experiment with 20-60 nuclei/cell type, mean ± SEM for each condition.

**Figure S9.**
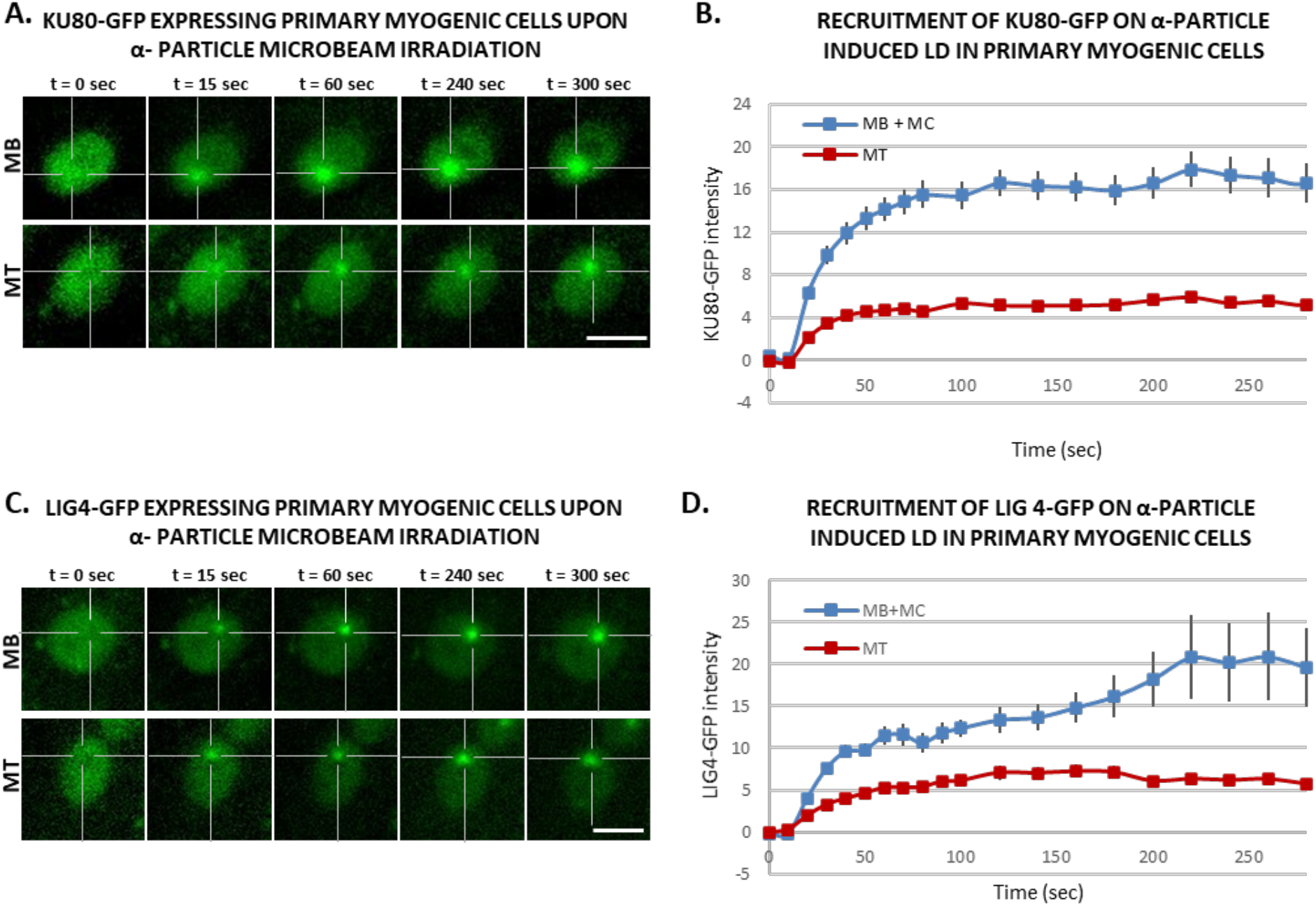
Activities of KU80 and LIG4 during myogenesis of primary cells. **A.** Sequential images of KU80-GFP recruitment to the locally damaged DNA (LD) by α-particle microbeam irradiation in transiently KU80-GFP expressing primary myoblasts (MB, upper panel) isolated from 4-6 days old mice and subsequent differentiated myotubes (MT, lower panel). The damaged areas are underlined by a dotted cross in the nucleus. Scale bar, 10 µm **B.** Recruitment curve of Ku80-GFP onto the LD by α-particle microbeam irradiation in primary myogenic cells isolated from 4-6days old mice transiently transfected with KU80-GFP expressing plasmid and then differentiated in myocytes (MB + MC, blue curve) and finally differentiated in MT (red curve). The irradiation was applied at t = 10 s. N ≥ 3 independent experiments with 42-65 nuclei/cell type, mean ± SEM. **C.** Sequential imaging of LIG4-GFP recruitment to the LD by α-particle microbeam irradiation in transiently LIG4-GFP expressing primary MB (upper panel) and MT (lower panel). The damaged areas are underlined by a dotted cross in the nucleus. Scale bar, 10 µm. **D**. Recruitment curve of LIG4-GFP on the LD by α-particle microbeam irradiation in transiently LIG4-GFP expressing primary MB + MC (blue curve) and MT (red curve). The irradiation was applied at t = 10 s. N ≥ 3 independent experiments with 24-32 nuclei/cell type, mean ± SEM.

**Figure S10.**
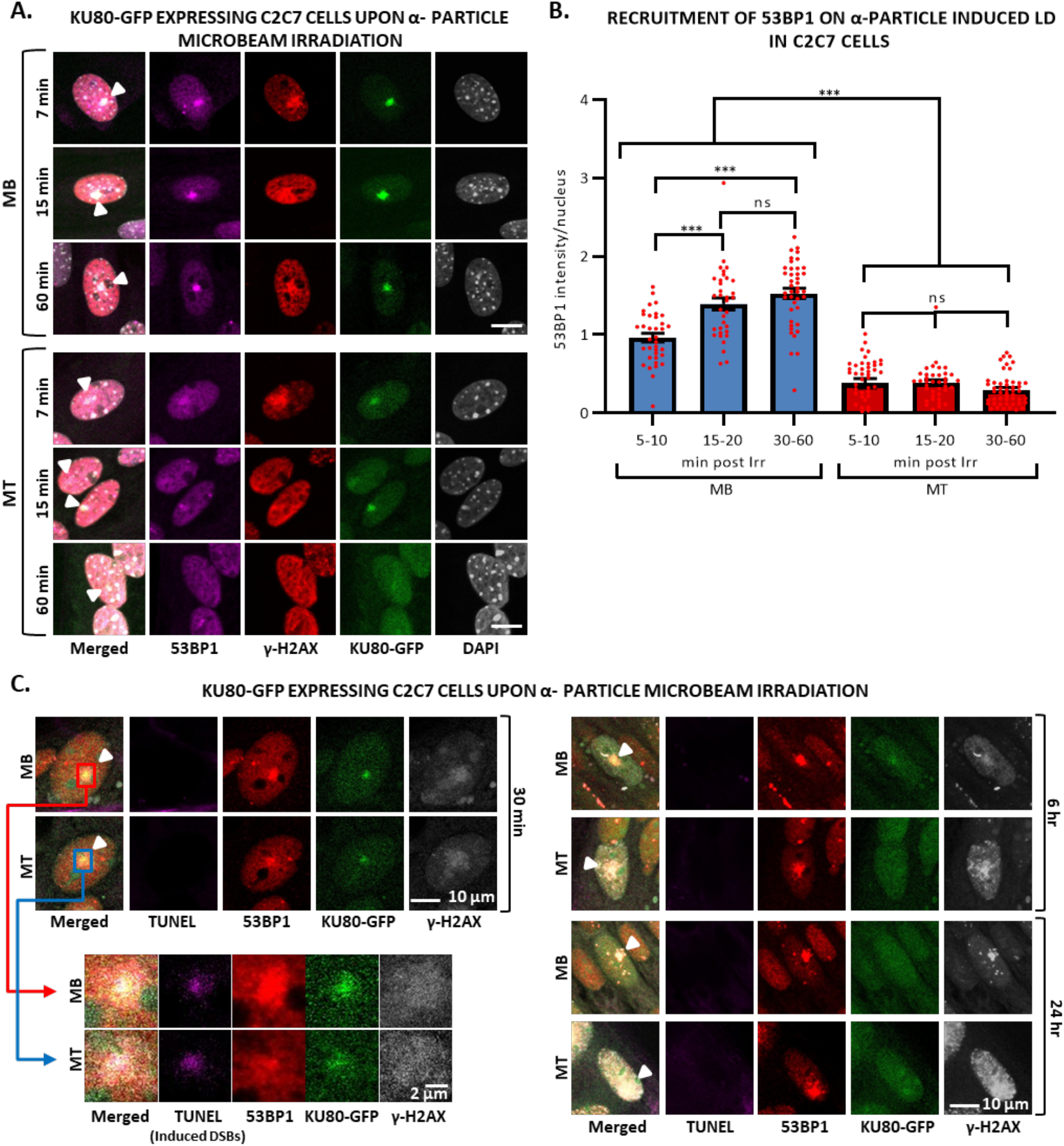
53BP1 starts to be visible in the late time-point in myotubes upon local DNA damage induction by α-particle microbeam irradiation. **A.** Representative images of stably KU80-GFP expressing C2C7 myoblasts (MB, upper panel) and myotubes (MT, lower panel) at the indicated time-point after locally damaging DNA (LD) by α-particle microbeam irradiation. Cells are immunolabelled with antibodies against DSB markers: 53BP1 (magenta) and γH2AX (red). Induced DNA damage site is indicated by a white arrowhead. DNA was stained with DAPI (grey). Scale bars, 10 µm. **B.** The level of 53BP1 recruited to the LD by α-particle microbeam irradiation represents the mean of maximum intensity upon immunofluorescence (IF) labelling with antibody against 53BP1 in stably KU80-GFP expressing C2C7 MB and MT from 5 to 60 min post-irradiation. Error bars represent the SEM obtained from at least 34-49 nuclei/time zone. Significance by 1-way ANOVA with post-hoc Tukey’s multiple comparison test, * p≤0.05, ** p≤0.01, *** p≤0.001. **C.** Representative images of stably KU80-GFP expressing C2C7 MB (upper panels) and MT (lower panels) at the indicated time-point after the LD by α-particle microbeam irradiation. Cells are immunolabelled with antibodies against DSB markers: 53BP1 (red) and γH2AX (white). TUNEL labelling is used to reveal DSBs (violet). Induced DNA damage site is indicated by a white arrowhead in the merged image. DNA was not stained. Scale bars, 10 µm. On the left part of the panel, the enlarged views are a zoom of the red and blue boxed regions and scale bars, 2 µm.

**Figure S11.**
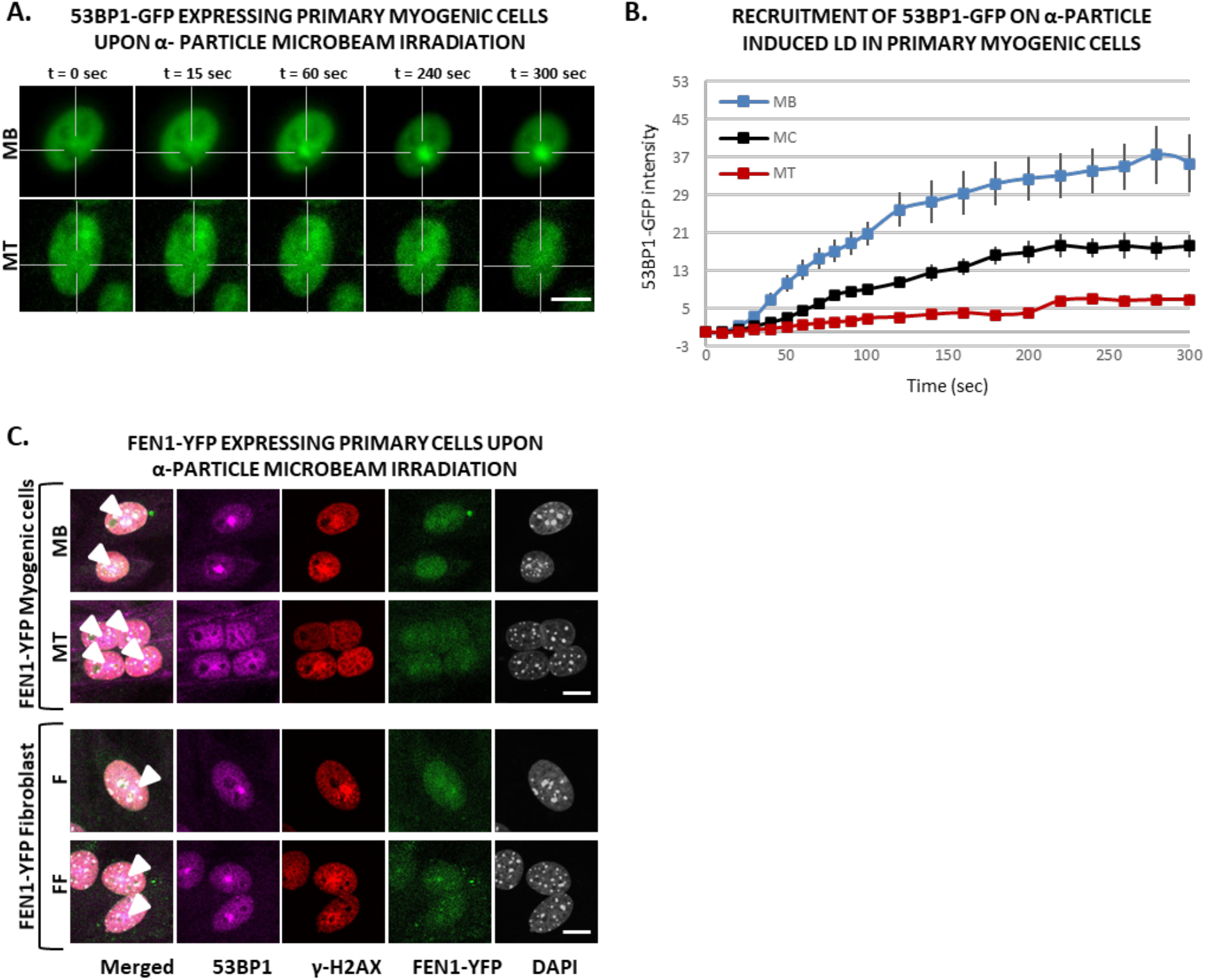
Decrease of 53BP1 recruitment upon α-particle microbeam irradiation in multi-nucleated cells is specific to myogenic cells. **A.** Sequential images of 53BP1-GFP recruitment to the locally damaged DNA (LD) by α-particle microbeam irradiation in transiently 53BP1-GFP expressing primary myoblasts (MB, upper panel) isolated from 4-6 days old mice and subsequent differentiated myotubes (MT, lower panel). The damaged areas are underlined by a dotted cross in the nucleus. Scale bar, 10 µm. **B.** Recruitment curve of 53BP1-GFP onto the LD by α-particle microbeam irradiation in transiently 53BP1-GFP transfected primary MB (blue curve), differentiating in myocytes (MC,black curve) and differentiated in MT (red curve). The irradiation was applied at t = 10 s. N ≥ 3 independent experiments with 19-26 nuclei/cell type, mean ± SEM. **C.** Representative images of primary MB and MT cells (upper panel), and mono-nuclear (F) and fused fibroblast (FF) (lower panel), isolated from FEN1-YFP mouse model, at the indicated time post α-particle microbeam irradiation. The cells are immunolabelled with antibodies against 53BP1 (violet) and γH2AX (red). DNA was stained with DAPI (grey), and local DNA damage site is marked with arrow-head in the merged images. Scale bar, 10 µm.

**Figure S12.**
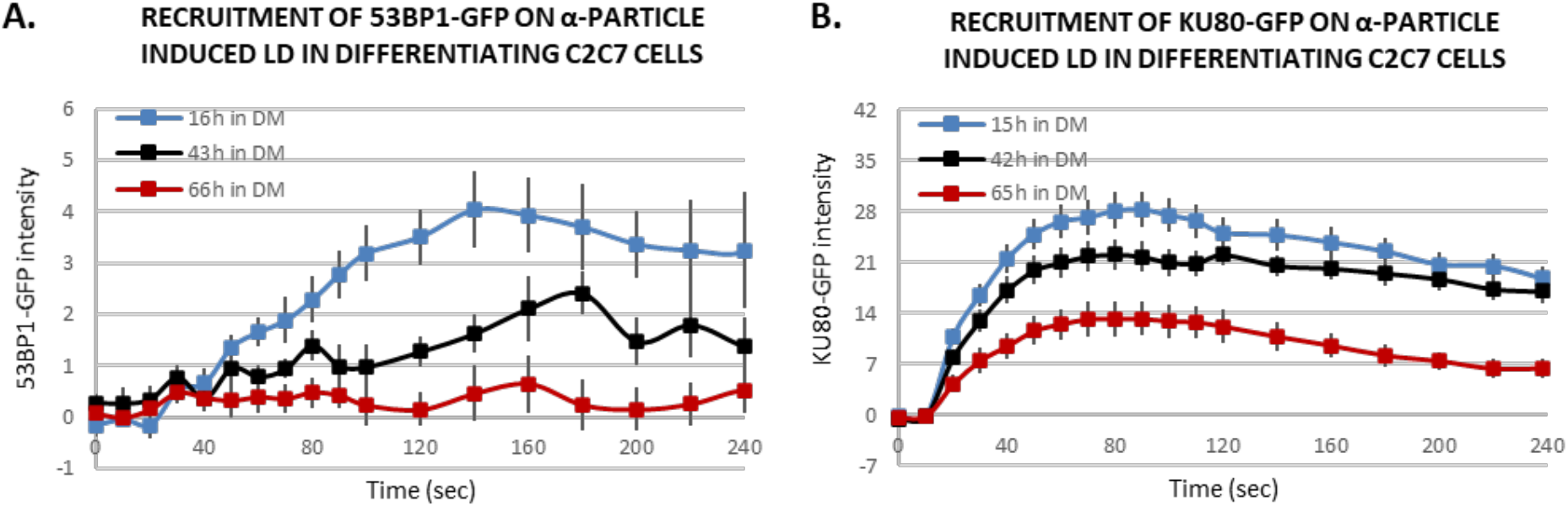
Decline of DNA damage response of 53BP1-GFP and KU80-GFP on α-particle microbeam irradiation -induced DNA damage sites along with differentiation. **A, B.** Recruitment curve of stably expressed 53BP1-GFP (A) and KU80-GFP (B) onto the locally damaged DNA (LD) by α-particles microbeam in stably transfected C2C7 cells in differentiation medium for 15-16h (blue), 42-43h (black) and 64-66 hours (red). The irradiation was applied at t = 10 s. mean ± SEM for each condition, 4-11 nuclei in (A) 15-26 cells in (B).

**Figure S13.**
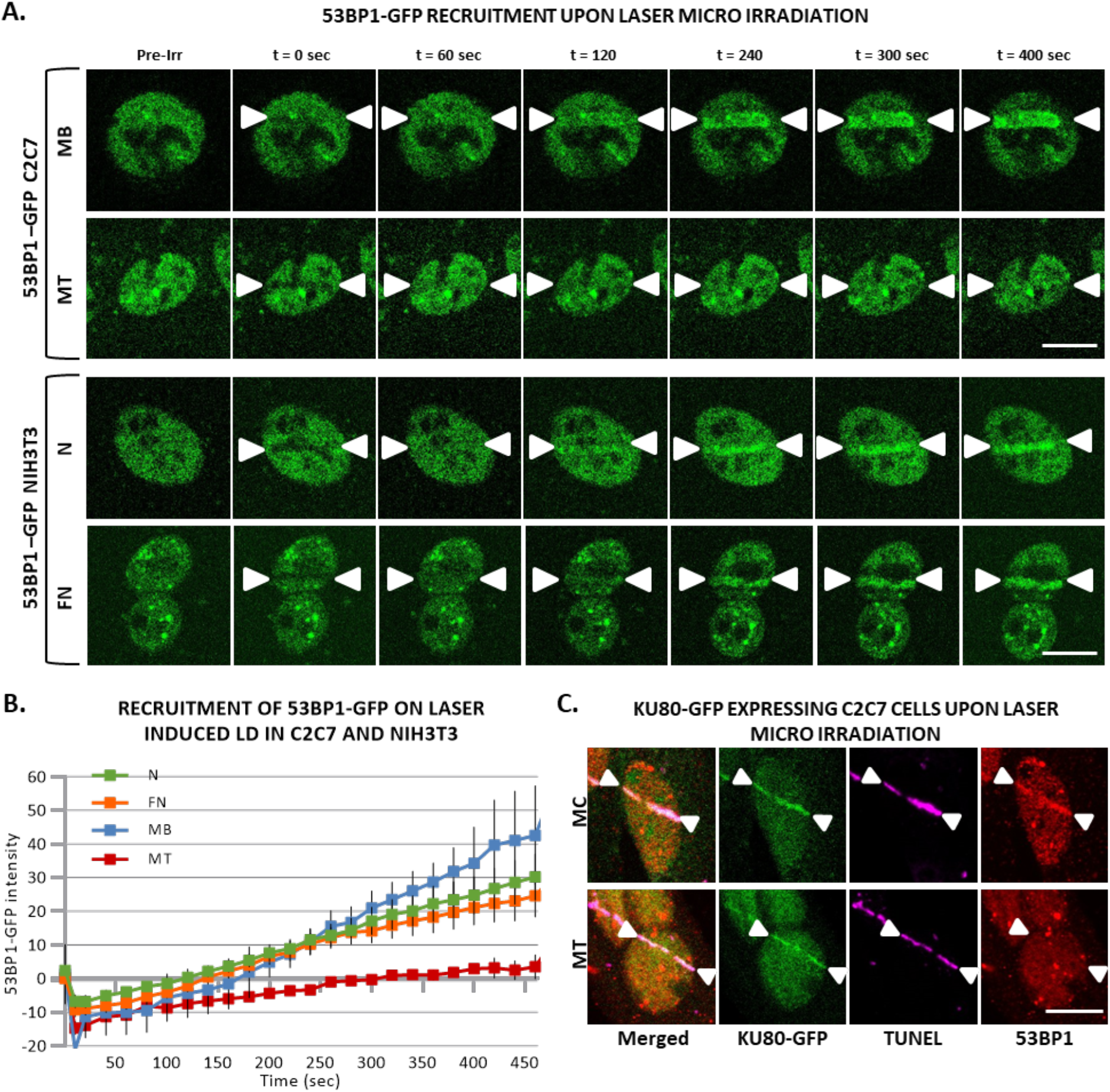
Almost no response of 53BP1 at local DNA damage sites induced by laser irradiation is specific to multinuclear myotubes. **A.** Sequential imaging of 53BP1-GFP recruitment to the locally damaged DNA (LD) by laser-irradiation in stably 53BP1-GFP expressing C2C7 myoblasts (MB) and myotubes (MT) (upper panel) or in transiently 53BP1-GFP expressing proliferative (N) and PEG-fused (FN) NIH-3T3 cells (lower panel). DNA damage site is indicated by 2 white arrow heads. Scale bar, 10 µm. **B.** Recruitment curve of 53BP1-GFP to the LD by laser-irradiation in stably 53BP1-GFP expressing C2C7 MB and MT (respectively, blue and red curves) or in transiently 53BP1-GFP expressing proliferative (N) and PEG-fused (FN) NIH-3T3 cells (respectively, green and orange curves). The irradiation was applied at t = 10 s. N ≥ 3 independent experiments with 20-34 nuclei/cell type), mean ± SEM. **C.** Representative images of stably KU80-GFP expressing C2C7 myocytes (MC, upper panel) and MT (lower panel) at the indicated time post-induced local DNA damage by laser-irradiation. Cells are immunolabelled with antibodies against the DSB markers 53BP1 (red). TUNEL labelling is used to reveal DSBs (violet). Induced DNA damage site is indicated by 2 white arrow heads. Scale bar, 10 µm.

